# Sustainable production of plastic-degrading enzymes in *Chlamydomonas pacifica*

**DOI:** 10.1101/2025.05.14.654053

**Authors:** Crisandra Jade Diaz, João Vitor Dutra Molino, Barbara Saucedo, Kalisa Kang, Évellin do Espirito Santo, Marissa Tessman, Abhishek Gupta, Michael D. Burkart, Ryan Simkovsky, Stephen Mayfield

## Abstract

The discovery of a new extremophile alga, *Chlamydomonas pacifica*, provides an opportunity to expand on heterologous protein expression beyond the traditional *Chlamydomonas reinhardtii. C. pacifica* is a unicellular extremophile capable of surviving at high pH, high temperatures, and high salinity. These various growth conditions allow C. pacifica to outcompete any invading contaminants in open-air environments. Developing this novel species as a platform for recombinant protein production could significantly advance commercial microalgal recombinant protein production. We have previously shown that *C. reinhardtii* can secrete a plastic-degrading enzyme: a PETase known as PHL7. This PETase is capable of cleaving ester bonds and has been used commercially for the degradation of PET plastics. However, the expression of such an enzyme has yet to be done in open raceway ponds and on a large scale. Here, we describe the culturing of PHL7 transgenic C. pacifica strain in three 80L raceway ponds and the measurements of recombinant enzymatic expression and activity found in the culture media. Our work provides proof of concept that this new organism can produce functional PHL7 enzymes in addition to producing the valuable components that inherently exist in the *C. pacifica* algae biomass.

**Graphical Abstract:** 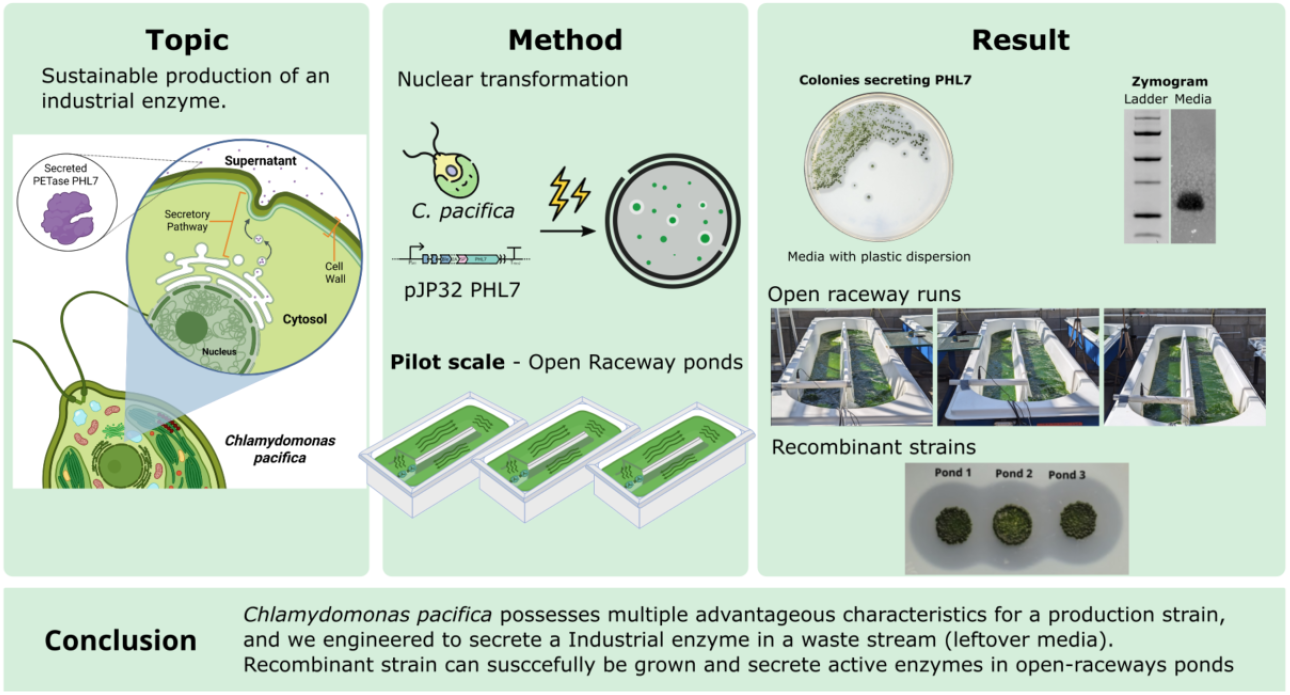

## Introduction

The rise in the number of studies highlighting the detrimental effects of microplastics has put a spotlight on the failings of current plastic management systems, especially plastic recycling (1–7). Traditional recycling consists of mechanical, chemical, and thermal recycling, each with its own caveats (8). However, despite increased calls to action for consumer recycling, a lack of recycling infrastructure as well as transparency in which products can actually be recycled, has led to very little actual recycling of plastics and a corresponding increase in the amount of plastic pollution. Only approximately 9% of plastics are recycled worldwide, and that number dwindles to a mere 5% in the United States (9). The remaining plastics are either burned or sent to landfills, or manage to make their way into the environment (10,11). Plastics have been found in every environment, from mountain tops to nearly every marine environment on the planet, there is nowhere to hide from plastic pollution (12,13). More recently, an even greater concern has developed over the discovery of microplastics and nanoplastics. These microscopic plastic particles are generated from the breakdown of larger plastic forms, and like their larger forms, persist in the environment due to resistance to biodegradation. Microplastics generally range from 5mm down to 1mm, and a newer subset known as nanoplastics ranges from 1 mm down to 1 μm (14). While the presence of micro- and nanoplastics has been reported in animals and food sources for some time, recent studies have begun to elucidate the various impacts that microplastics have on human health (7). Microplastics are now associated with heart disease and stroke, lowered sperm count, and have been found to cross the blood-brain barrier, and also can pass between mother and fetus via the placenta (2,4,6). Due to their persistence and ubiquitous presence, we can only expect to see more studies showing the relationship between microplastics and human health issues.

One of the few ways to combat plastic pollution and the creation of persistent microplastics is through biological or enzymatic degradation. Biological degradation is one facet of the biodegradation process whereby organisms can produce enzymes capable of breaking down the bonds within the plastics (15,16). For a product to be considered biodegradable, the organism responsible for the biological degradation must also use individual plastic monomers as an energy source, usually in the form of carbon. However, characterization of individual enzymes capable of breaking these bonds creates another platform for addressing plastic pollution. Enzymes involved in biological degradation can be isolated and produced in an industrial platform (10). Degrading plastics with enzymes down to their individual monomers allows these monomers to potentially be upcycled into new plastic products without compromising the structural integrity (unlike current recycling methods) (17,18). One such enzyme is a PETase known as PHL7. PHL7 was discovered in a metagenomic dataset from compost and was found to be an extremely robust enzyme (19). PETases are enzymes found to be capable of degrading PET plastics. This is done by hydrolyzing the ester bonds within the PET polymer. PETases fall under the category of “Esterase” enzymes, although not all esterases are capable of PET degradation (20,21).

Here we have chosen to express *PHL7* in the new extremophile strain known as *Chlamydomonas pacifica* (22). This extremophile can grow in high salt, pH, and temperatures up to 42°C. *C. pacifica* was first identified during a bioprospecting excursion for unicellular algae along the California coast. Ironically, it was discovered on campus at the University of California, San Diego. While highly similar in morphology to *C. reinhardtii*, the genetic relationship between the two species is more distant compared to Volvox sp., which have very different morphologies (22). Regardless, *C. pacifica* still lies within the clade of *Reinhardtinia* and has enough genetic similarity to the *C. reinhardtii* genome for genetic tools developed for *C. reinhardtii* to be directly transferred to this new strain. *Chlamydomonas* species are especially well-placed for their use as a platform for protein expression (23). Various studies support the successful production of recombinant proteins and other natural products from *Chlamydomonas reinhardtii*. While there is a definite genetic distinction, our lab has found that the tools developed for this model organism are translatable to this new extremophile strain (24,25).

*Chlamydomonas* is a versatile genus of green microalgae with significant potential for products in food, oil, and biodegradable biopolymer industries (24,26,27). In *Chlamydomonas reinhardtii*, the production of recombinant proteins has become a cornerstone of algal biotechnology, offering a versatile platform for generating a wide range of bioactive molecules (28–31). One of the key advantages of using *C. reinhardtii* is its ability to efficiently secrete recombinant proteins into the culture medium, which simplifies the purification process and reduces costs (25,32). Secretion of proteins is particularly important because it enables the development of methods for the direct harvesting of target proteins from the growth medium, avoiding the complex and expensive steps of cell disruption and intracellular protein extraction (33). This feature has the ability to enhance the scalability of production and lower the overall cost of protein recovery. Moreover, the secreted proteins are often correctly folded and functionally active, making *C. reinhardtii* an attractive system for producing therapeutic proteins, industrial enzymes, and other valuable bioproducts (25). The ability to harness *C. reinhardtii* for large-scale protein production underscores its potential in both academic research and industrial applications, making it a vital tool in the field of biotechnology (34).

The extremophilic properties of *Chlamydomonas pacifica*, particularly its adaptation to high-salinity environments, offer distinct advantages in biotechnological applications. Unlike many freshwater algae, *C. pacifica* thrives under extreme conditions, such as elevated salinity, which can lead to the production of specialized enzymes and metabolites with unique properties (22). These extremophilic traits enable *C. pacifica* to produce enzymes that remain active and stable in harsh environments, making them valuable for industrial processes that require robustness and resilience (25). Additionally, the ability to grow in saline environments allows for the utilization of non-freshwater resources, potentially reducing the competition for agricultural water supplies and promoting sustainable practices. Not only does this give *C. pacifica* an advantage over other algal growth platforms, but can also compete with other agricultural crops. Most traditional crops are not well-suited to grow in brackish water, a largely untapped resource in the US, underlying the country with more than 800 times the amount currently used yearly (35).

## Results

### Expression and Secretion of the PET-Degrading Enzyme PHL7 in *C. pacifica*

To evaluate the feasibility of expressing a plastic-degrading enzyme in *C. pacifica*, a synthetic gene encoding the PETase PHL7 was codon-optimized for nuclear expression and inserted into the vector pJP32 under the control of the strong, constitutive *PAR1* promoter (**Figure 1A**). To enable secretion of the enzyme, the N-terminal region of PHL7 was fused to SP7, a secretion signal peptide derived from the *C. reinhardtii* cell wall protein SAD1p (32). Additionally, the construct included a bleomycin resistance gene (*ble*) for selection, separated by a self-cleaving 2A peptide (F2A), allowing co-expression of both genes from a single transcriptional unit. The transgene cassette was terminated with the *rbcs2* 3′-UTR to enhance mRNA stability.

**Figure 1.**
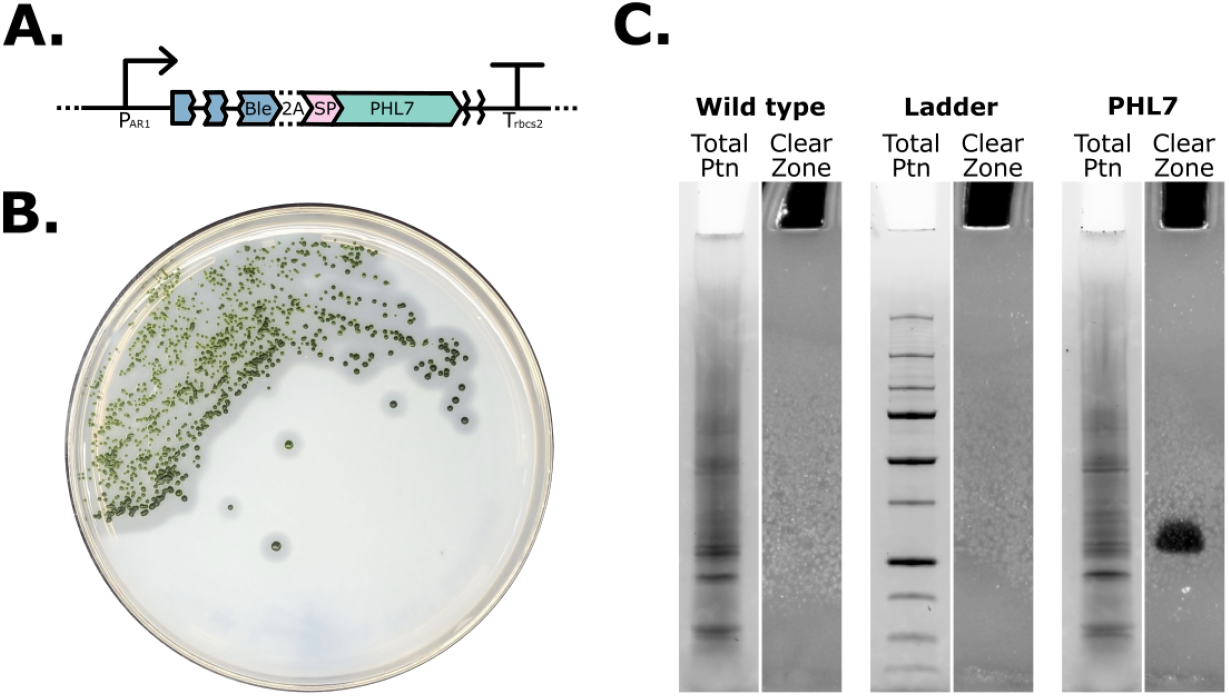
Expression and Activity of the Plastic-Degrading Enzyme PHL7 in *C. pacifica*. **(A)** Vector map of the *pJP32 PHL7* construct, showing the expression elements for the plastic-degrading enzyme PHL7 under the control of the PAR1 promoter, including a Bleomycin resistance gene (Ble), a Food and Mouth 2A self-cleaving peptide (2A), and the signal peptide from XXX (SP) for proper secretion. **(B)** *C. pacifica* grown on TAP/Impranil® DLN agar plate, displaying clear halos around colonies. These halos indicate enzymatic degradation of the polymer, suggesting active plastic-degrading enzyme secretion by *C. pacifica*. **(C)** Zymogram with Impranil-DLN as a substrate. The total protein (protein auto fluorescence in Stain Free method) and clearing zone (amido black method) are shown for both the wild-type and PHL7-transformed *C. pacifica* supernatants. Only the supernatant of *C. pacifica* expressing PHL7 exhibits a clearing zone, indicating degradation of the polymer by the plastic-degrading enzyme.

Following transformation of wild-type *C. pacifica*, stable nuclear transformants harboring the PHL7 expression construct (pJP32PHL7) were isolated based on antibiotic resistance. To assess functional secretion of the recombinant enzyme, transformants were screened using a plate-based halo assay with Impranil DLN, a polyester-polyurethane that forms a turbid dispersion in agar and serves as a model substrate for polymeric ester bond hydrolysis. As shown in **Figure 1B**, colonies expressing PHL7 produced distinct clearing zones around the colony perimeter, indicating localized polymer degradation in the surrounding medium. In contrast, wild-type *C. pacifica* colonies did not exhibit halo formation under identical conditions, confirming that degradation was due to recombinant enzyme activity (**Supplementary Figure 1**).

To further validate enzyme secretion and activity, culture supernatants were analyzed using an in-gel activity assay (zymography) in which SDS-PAGE was performed on gels co-polymerized with Impranil as a substrate. Only the supernatant from PHL7-expressing transformants exhibited a clear lytic band indicative of enzymatic degradation, while the wild-type supernatant showed no detectable activity (**Figure 1C**). This result demonstrates that PHL7 is not only successfully expressed and secreted by *C. pacifica* but also remains catalytically active in the extracellular environment.

### Laboratory-Scale Growth and PET Degradation by *C. pacifica* Expressing PHL7

To determine whether recombinant expression of PHL7 imposes any fitness cost on *Chlamydomonas pacifica*, we monitored the growth kinetics of PHL7-expressing transformants in comparison to the wild-type strain under laboratory-scale culture conditions. Both strains were grown in parallel in HA medium in batch shake-flask cultures, and optical density at 750 nm (OD_750_) was measured over a 7-day period. As shown in **Figure 2A**, the growth profile of *C. pacifica/PHL7* closely mirrored that of the wild type, with no evidence of impaired growth. In fact, PHL7-expressing cells exhibited a marginally higher OD during the early exponential phase, although both strains reached similar stationary-phase densities. These data indicate that integration and expression of the PHL7 construct do not negatively affect the host’s proliferation under standard culture conditions.

**Figure 2.**
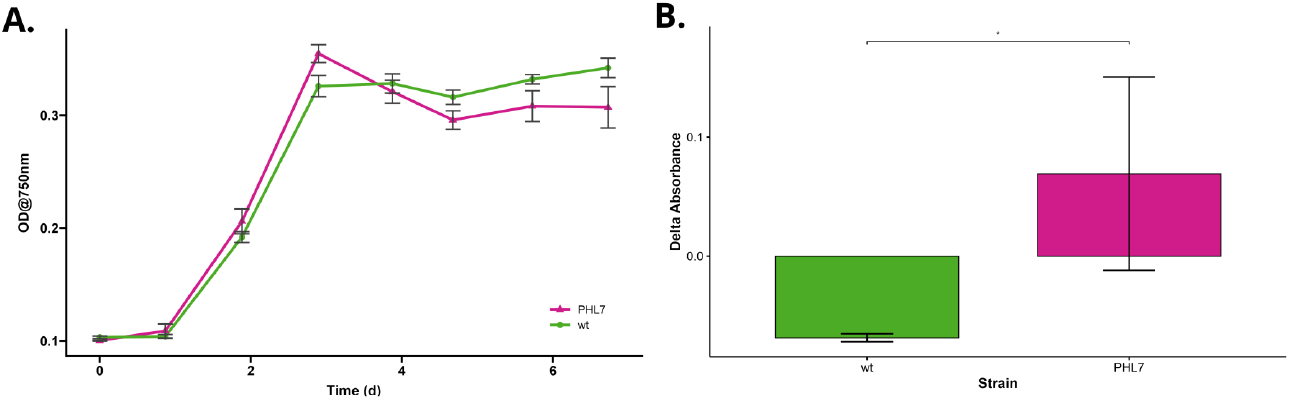
Figure 2A: Growth curve comparison between wildtype (WT) *C. pacifica* and *C. pacifica/PHL7* grown in parallel at lab scale. Figure 2B: Absorbance at 240 nm of supernatants incubated with PET beads. The supernatant containing PHL7 had a statistically significant increase in the change in absorbance at 240 nm compared to WT. *C. pacifica*, indicating a higher level of TPA release. Figure 3B: Zymogram indicating successful secretion of PHL7 and its activity compared to WT.

To assess the enzymatic activity of secreted PHL7 against polyethylene terephthalate (PET), culture supernatants from *C. pacifica/PHL7* and wild-type controls were incubated with PET beads (50% crystallinity) at 70 °C for 7 days. PET depolymerization was evaluated by measuring the release of terephthalic acid (TPA), a primary degradation product, via absorbance at 240 nm (25). Supernatants from PHL7-expressing strains exhibited a significantly greater increase (*p-value* = 0.04255) in A_240_ compared to wild-type controls (**Figure 2B)**, consistent with enzymatic PET hydrolysis. Collectively, these results confirm that recombinant PHL7 is both expressed and secreted in a catalytically active form by *C. pacifica* and that it retains the ability to depolymerize PET into its monomeric components without adversely impacting host growth.

### Scale-Up and Greenhouse Performance of *C. pacifica/PHL7* in Open Raceway Ponds

To evaluate the scalability and robustness of recombinant PHL7 production under conditions relevant to industrial biotechnology, the engineered *C. pacifica/PHL7* strain was cultivated in three replicate 80-liter open raceway ponds housed within a greenhouse facility. Each pond was inoculated with pre-adapted culture from carboy-scale pre-cultures (**see Video 1)** and operated in parallel over a 12-day period (**Figure 3A**). The ponds were mixed continuously using propeller agitation, and environmental parameters—including photosynthetically active radiation (PAR), temperature, and pH—were monitored in real time using embedded sensors.

**Figure 3.**
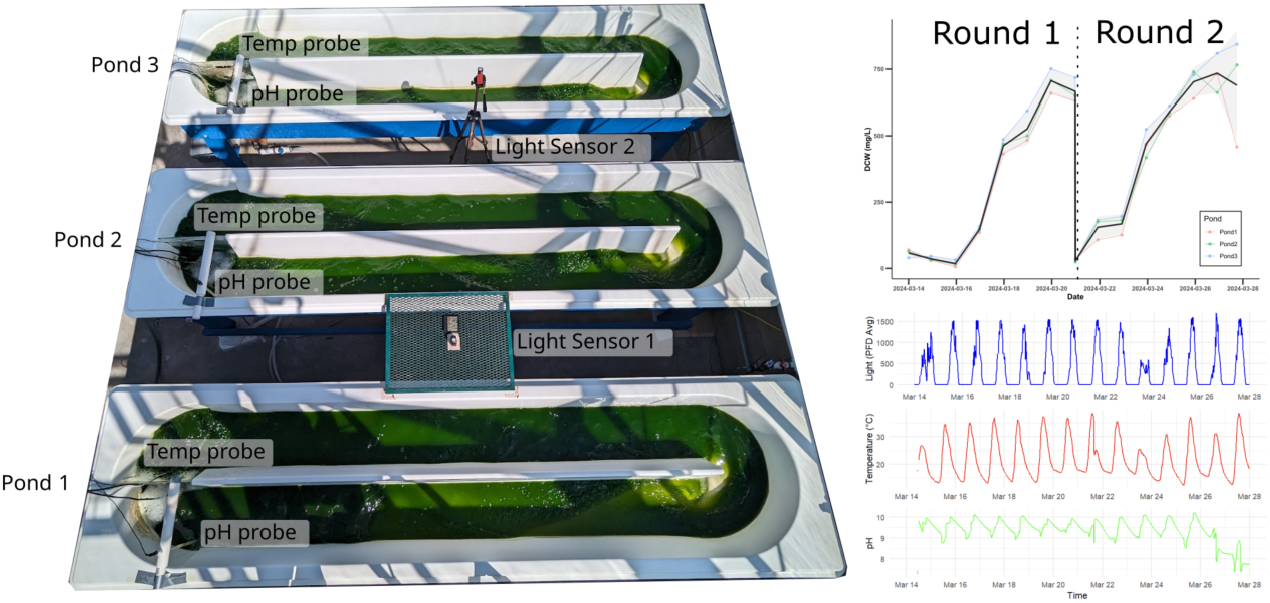
Open raceway ponds culturing. Figure 3A: *C. pacifica/PHL7* grown in parallel in 80L raceway ponds. All three ponds were biological replicates that were simultaneously measured for light intensity, temperature, and pH over the course of 12 days. Figure 3B: Dry cell weight (DCW) was measured daily for each pond. Figure 3C: Sensor data for light, temperature, and pH over time.

Dry cell weight (DCW) measurements indicated consistent biomass accumulation across all three ponds, with cultures reaching peak densities of approximately 1.0 g/L by the end of the cultivation period (**Figure 3B**). Following a brief lag phase during the initial adaptation (days 0–3), cultures entered exponential growth and exhibited comparable performance across replicates. The slight delay in early growth was likely due to physiological acclimation from controlled indoor culture to the dynamic conditions of the raceway environment.

Environmental conditions varied according to diurnal and weather-dependent fluctuations. Light availability followed the natural daylight cycle, with substantial variability in daily light integral (DLI) due to intermittent cloud cover (**Supplementary Figure 2**). Growth rates correlated positively with light intensity, underscoring the photoautotrophic nature of *C. pacifica* and its tolerance to high light intensity in peak hours per day. Temperature in the ponds exhibited a diel oscillation from approximately 14 °C during nighttime to over 37 °C during peak daytime (**Figure 3C**, middle panel), mirroring the ambient greenhouse climate. The strain tolerated this range without visible signs of stress, consistent with its thermotolerant properties (22).

Throughout the cultivation, pond pH was manually regulated and maintained near 10.5 (**Figure 3C**, bottom panel), a condition known to confer a selective advantage to *C. pacifica* by inhibiting bacterial and eukaryotic contaminants. pH adjustment was performed daily to sustain alkaline conditions within the optimal physiological range for the host.

These results confirm that the engineered *C. pacifica/PHL7* strain remains physiologically robust under large-scale, non-sterile, and environmentally variable conditions. Crucially, the strain maintained healthy growth and biomass productivity in open systems while sustaining the expression of a functional heterologous enzyme. This highlights *C. pacifica*’s potential as a viable chassis for outdoor biotechnological applications, including plastic degradation.

### Enzyme Activity Retention Following Raceway Pond Cultivation

To assess whether the plastic-degrading enzyme PHL7 remained expressed and catalytically active after outdoor-scale cultivation, samples from all three raceway ponds were collected at the end of each 6-day run and analyzed using multiple activity assays. Cells and supernatants were harvested from each pond and replicated at the final time point of two independent cultivation trials (Rounds 1 and 2). They were then tested for enzyme activity using zymography and quantitative polymer degradation analysis.

To confirm enzymatic activity in pond cultures, supernatants were concentrated approximately 400-fold and subjected to SDS-PAGE-based zymography using Impranil as a copolymerized substrate. Clear degradation bands were observed in all pond samples (**Figure 4A, B**), providing strong evidence that functional PHL7 accumulated extracellularly during pond cultivation. Notably, enzymatic activity was retained despite the challenging physicochemical conditions of the open system, including elevated pH and fluctuating temperatures.

**Figure 4.**
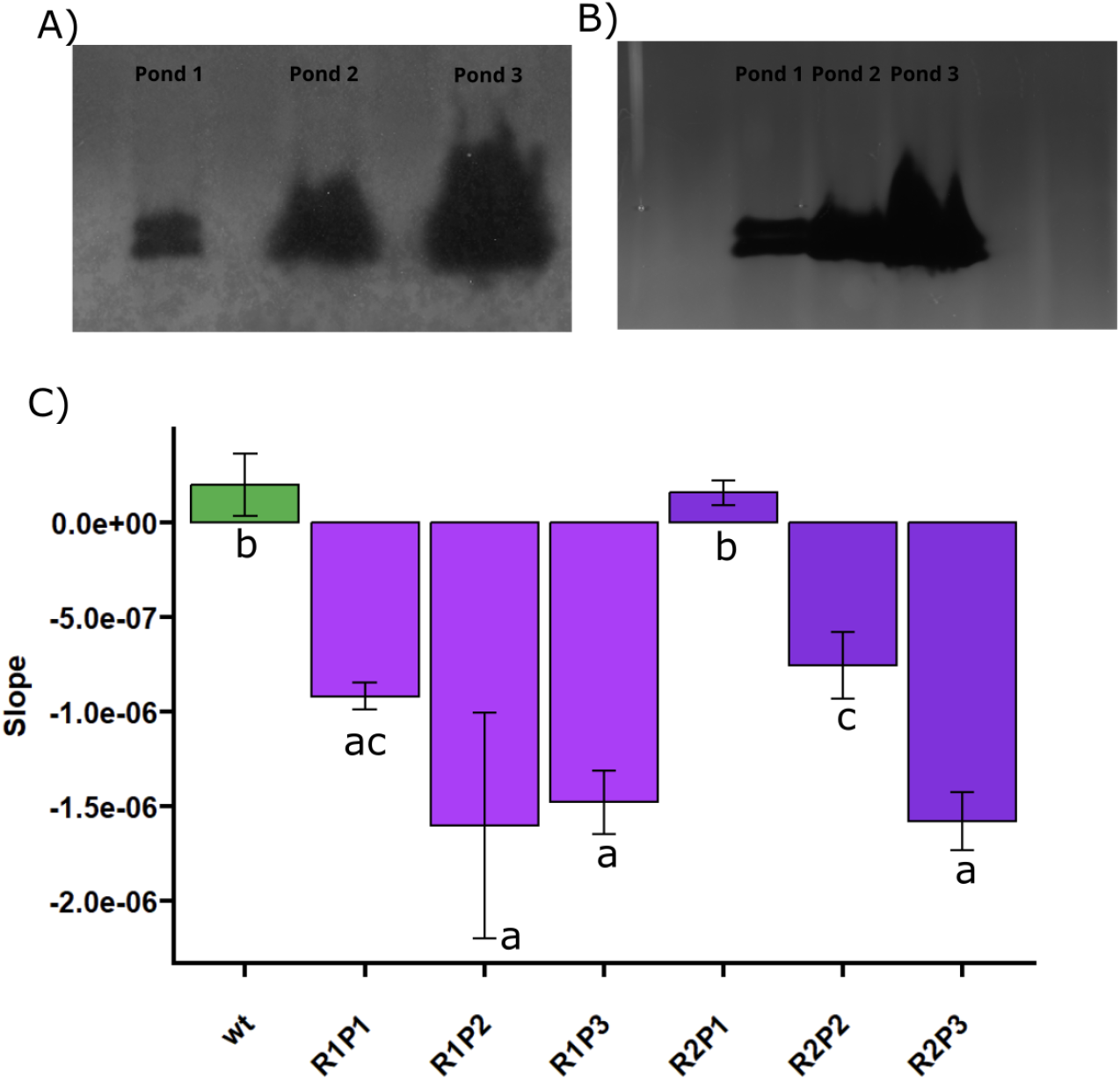
Impranil degradation by genetically engineered microalga *Chlamydomonas pacifica* expressing PHL7 in open-pond systems. **(A–B)** Visual documentation of Impranil degradation of supernatant samples from the three replicate open ponds (Pond 1–3) at pre-harvesting of genetically modified *Chlamydomonas pacifica* strains harboring the plasmid **PJP32PHL7**. Ponds were incubated outdoors for two consecutive 7-day rounds. **(C)** Quantitative analysis of polymer degradation, expressed as the slope of absorbance (ΔAbs/time) over the incubation period. A more negative slope corresponds to faster degradation of Impranil and higher enzymatic activity. Each bar represents the mean ± standard deviation of biological replicates for wild-type *C. pacifica* (wt) and engineered strains from Round 1 (R1P1–R1P3) and Round 2 (R2P1–R2P3). Distinct lowercase letters indicate statistically significant differences between strains (Tukey’s HSD post hoc test following one-way ANOVA, *p* < 0.05).

Because PHL7 exhibits optimal catalytic performance at near-neutral to mildly alkaline pH, a pH shift was implemented in the second cultivation trial (Round 2) to examine whether enzyme yield could be improved. Specifically, ponds were maintained at pH 10.5 throughout most of the experiment but adjusted to pH 8.0 during the final 48 hours before harvest to favor de novo enzyme synthesis under more favorable catalytic conditions. However, no consistent enhancement in zymogram signal intensity or halo formation was observed relative to Round 1, which was conducted entirely at pH 10.5. These results suggest that PHL7 is sufficiently stable and active under high-pH conditions and that pH reduction prior to harvest does not substantially impact active enzyme accumulation.

Quantitative assessment of polymer degradation was conducted by measuring the rate of Impranil breakdown, expressed as the slope of absorbance change over time (ΔAbs/time). As shown in **Figure 4C**, the majority of PHL7-expressing pond cultures (R1P1–R1P3 and R2P2–R2P3) exhibited significantly more negative slopes compared to the wild-type control, consistent with high levels of enzymatic activity. An exception was observed for R2P1, which showed markedly reduced activity. This reduction coincided with signs of biomass decline evident in the growth data (**Figure 3**), potentially linked to both the artificial pH reduction implemented during the final 48 hours of cultivation and lower light availability for that pond. Despite this outlier, most engineered strains demonstrated consistent and statistically significant Impranil degradation relative to the wild type (ANOVA with Tukey’s HSD, *p* < 0.05). No overall differences in enzyme activity were detected between Round 1 (pH 10.5 throughout) and Round 2 (pH shift to 8.0 at the end), further supporting the conclusion that high-pH outdoor cultivation does not impair PHL7 expression or enzymatic function.

Overall, these data demonstrate that *C. pacifica/PHL7* maintains stable transgene expression and secretes catalytically active PETase under outdoor, non-sterile cultivation conditions, with enzyme activity detectable directly from pond supernatants.

## Discussion

### Feasibility of *C. pacifica* as a Platform for Enzyme Production

Our results demonstrate *Chlamydomonas pacifica* as a viable new platform for the sustainable production of recombinant plastic-degrading enzymes. The successful expression and secretion of the PET hydrolase PHL7 in *C. pacifica* (**Figure 1**) shows that genetic tools developed in *C. reinhardtii* can be directly transferred to this extremophilic alga. The use of a signal peptide (SP7) enabled efficient secretion of PHL7 into the culture medium, consistent with previous demonstrations of secretory expression in *C. reinhardtii* (32). Notably, the PHL7-expressing *C. pacifica* grew as robustly as the wild-type strain (**Figure 2A**), indicating that the metabolic burden of producing and secreting this enzyme did not noticeably hinder growth. This is an encouraging result, as in some hosts the overexpression of heterologous enzymes can reduce fitness or yield (36). The lack of growth penalty in *C. pacifica* may be attributed to the stability and folding efficiency of PHL7, as well as the strong growth characteristics of *C. pacifica* itself. The *C. pacifica/PHL7* strain was able to withstand large-scale open pond cultivation (80 L) under non-sterile, variable environmental conditions, maintaining both its growth rate and protein secretion phenotype (**Figures 3 and 4**). This robustness underscores one major advantage of *C. pacifica* as a production host: its extremophilic tolerance (e.g., high pH, high salt) allows culture in open systems with minimal contamination, a feat not easily achievable with conventional hosts like *E. coli* or yeast and other algae (37). Previous studies have noted the feasibility of outdoor or greenhouse cultivation of *Chlamydomonas* pacifica for biotechnology applications (24). Our work extends this to recombinant enzyme production, demonstrating that an active enzyme can be produced in situ at pilot scale.

Compared to conventional microbial platforms, *Chlamydomonas pacifica* offers a compelling combination of sustainability, operational simplicity, and economic efficiency. While heterotrophic hosts such as *Escherichia coli* and *Saccharomyces cerevisiae* are widely used for recombinant protein production due to their rapid growth and high protein yields, these systems typically require sterile fermenters, refined carbon sources, and strict environmental controls—factors that increase both complexity and cost. In contrast, *C. pacifica* is a photoautotrophic microalga capable of growth in open raceway ponds using only sunlight, CO2, and inorganic nutrients, thereby substantially reducing input requirements.

A further advantage of *C. pacifica* lies in its potential to support integrated bioprocessing. As demonstrated previously (24), this strain can serve as a source of multiple value-added products, including lipids for biofuel, starch for bioplastics, and secreted proteins from the culture supernatant. The ability to extract these components from distinct cellular or extracellular fractions opens opportunities for multi-product biorefineries and enhances economic viability.

In our raceway pond experiments, *C. pacifica* cultures derived energy exclusively from natural light and were maintained at a high pH (∼10.5), which inherently suppresses microbial contaminants—an environmental condition unsuitable for most traditional heterotrophs. This extremophilic cultivation strategy functions as a passive biocontainment system, supporting non-sterile outdoor production while minimizing ecological risk. The success of our 12-day pond trial underscores the practicality of this approach in real-world biotechnological applications.

Crucially, the recombinant PET-degrading enzyme PHL7 remained catalytically active throughout outdoor cultivation, despite prolonged exposure to alkaline conditions and fluctuating temperatures ranging from 14 °C to 37 °C. Enzymatic activity was also preserved after the culture medium was adjusted to a more moderate pH prior to harvest (**Figure 4D**), indicating that PHL7 is resilient under shifting environmental stress. Unlike many enzymes that irreversibly denature outside their optimal conditions, PHL7 retained functional integrity, likely reflecting its origin as a thermostable polyester hydrolase (19,38). This robustness supports its application in environmental plastic degradation. Further, it underscores the suitability of *C. pacifica* as a production host for other extremozymes—enzymes evolved to function in harsh or variable conditions.

In parallel, we explored a bioprocessing strategy to improve the recovery of potentially more sensitive recombinant proteins by shifting the pond pH to a more neutral value during the final 48 hours of cultivation. The rationale was to maintain high pH during the growth phase, favoring contamination resistance and algal proliferation, while creating a more favorable environment for recombinant protein folding and stability during the production phase. Although this pH-shift approach was not essential for PHL7 recovery, it may prove beneficial for future applications involving less stable heterologous proteins.

In our trial, the recovery of enzymatic activity varied across pond replicates (**Figure 4C**). Notably, one pond (R2P1) exhibited markedly lower activity despite undergoing the same pH-shift protocol as the other ponds. Analysis of growth and environmental sensor data revealed that Pond 1 consistently experienced reduced light exposure due to intermittent shading, partially captured by the light sensors positioned near Pond 2. These findings suggest that both environmental microconditions (e.g., light availability) and process variables (e.g., pH regime) influence recombinant protein yield and stability in open-pond systems. While the pH-shift strategy may be useful for enhancing the recovery of sensitive proteins, further optimization, including a light management plan and real-time physiological monitoring, will be required to maximize its effectiveness.

Despite the advantages of using *C. pacifica* as a production platform, certain limitations persist in the current system. A notable constraint is the relatively low titer of secreted PHL7 enzyme in the culture supernatant. Although enzymatic activity was detectable after approximately 400-fold concentration of the pond supernatant (**Figure 4C**), this indicates a modest enzyme concentration in the bulk culture. Specifically, in Pond 1 during Round 2 at harvest, the PHL7 concentration was estimated at 24 μg/L, which is suboptimal for a single-product process.

However, this yield can be enhanced through various strategies. To date, no specific optimizations have been implemented concerning the process, strain, or vector design. In *Chlamydomonas reinhardtii*, several successful approaches have been demonstrated that could be adapted to *C. pacifica*. For instance, the addition of glycomodules to recombinant proteins has been shown to improve secretion efficiency (39). Moreover, optimizing culture media and harvest strategies can significantly influence protein yields (40,41). Breeding techniques, such as mating and selection for strains with enhanced expression capabilities, have also been explored in *C. reinhardtii* (42).

Low yields of secreted heterologous proteins are a known challenge in *Chlamydomonas* species, often attributed to bottlenecks in the secretory pathway (43). Future improvements could involve optimizing genetic elements, including promoters, codon usage, and secretion signals. Engineering the host strain by knocking out proteases or modifying components of the secretory pathway may also enhance protein stability and secretion (44). Notably, previous studies in *C. reinhardtii* have screened multiple signal peptides to enhance secretory expression (32), and similar strategies could be applied to *C. pacifica*.

Quantifying the actual enzyme concentration in the culture, using methods such as ELISA, would provide benchmarks against other production platforms. For example, PHL7 and related PET hydrolases have been produced in *Escherichia coli* and purified to high concentrations suitable for industrial applications (17). Expression systems like *Pichia pastoris* often yield tens to hundreds of milligrams per liter of secreted enzyme (45). While *C. pacifica* may not yet achieve these titers, the cost savings and sustainability of algae-based production could offset the need for very high yields in specific applications.

Additionally, the enzyme produced by *C. pacifica* is secreted directly into the extracellular medium, simplifying downstream processing. This could potentially eliminate the need for costly purification steps if the crude supernatant is used directly for plastic waste treatment. Furthermore, co-products retrieved from the biomass, such as biofuel from lipids, bioplastics from starch, and animal feed from proteins, could provide additional revenue streams, enhancing the overall economic viability of the bioprocess (24).

### Enzymatic Plastic Degradation in the Context of Prior Work

The functionality of the algae-produced PHL7 was confirmed through PET plastic degradation assays (**Figure 2B**), which showed the release of TPA, a signature monomer of PET hydrolysis. The levels of TPA released by *C. pacifica* supernatant were modest but significant, reflecting the activity of the enzyme without any purification or enhancement. It is instructive to compare this outcome to other PET-degrading enzyme systems. PHL7 is one of the more efficient PET hydrolases reported in the literature: for example, purified PHL7 can **completely depolymerize** low-crystallinity PET films, releasing on the order of 90 mg/g PET of terephthalate within 1–2 days under optimal conditions (19). An engineered variant of a related enzyme has achieved even faster depolymerization – Tournier et al. (2020) report 90% conversion of PET to monomers in just 10 hours using a high-performance cutinase mutant at 72 °C (17). Compared to these controlled reactions with purified enzymes, the PET degradation we observed with algal culture supernatant is lower, likely due to the lower enzyme concentration and the use of a crude preparation. Nevertheless, the fact that any PET degradation occurred directly in the culture medium is promising. It demonstrates that the secreted PHL7 retained its catalytic function and appropriate folding. By performing activity assays at 70 °C, we took advantage of PHL7’s thermostability; many mesophilic PETases (such as the original *I. sakaiensis* PETase) would be inactivated at this temperature, whereas PHL7 remains active (46). This highlights a benefit of our chosen enzyme: its thermal optimum aligns well with conditions needed to soften PET substrates (near PET’s glass transition temperature), thereby enhancing overall depolymerization efficiency (17). In the context of enzymatic recycling, using a thermophilic enzyme like PHL7 can significantly improve reaction rates and yields. Our work is the first to produce PHL7 in a photosynthetic extremophilic organism, and the successful activity assays confirm that *C. pacifica* can serve as a biofactory for enzymes that act on persistent plastics.

### Implications and Future Directions

The present work demonstrates a proof-of-concept for sustainable enzyme production in a microalgal system, with direct application to plastic waste mitigation. To our knowledge, this is the first report of a PETase enzyme produced in algal open ponds and shown to actively depolymerize plastic materials. The ability to grow engineered *C. pacifica* in non-sterile conditions at scale is a significant step toward economically viable biomanufacturing. In contrast to heterotrophic expression systems that rely on expensive inputs (sugars, organic growth media) and tight environmental control, our algae-based approach leverages free resources (sunlight, CO2, saline water) and the natural resilience of an extremophile species. This strategy markedly reduces the carbon footprint and energy demands of enzyme production. We envision that *C. pacifica* cultures could be integrated into a circular plastic management workflow: the algae would produce enzymes that break down plastic waste into monomers, and simultaneously, the algal biomass could be harvested to serve as raw material for new bioplastics or other valuable bioproducts. Indeed, *C. pacifica* is known to accumulate lipids and starch under certain conditions (24), which could be extracted for biofuel or biopolymer production (e.g., polylactic acid, polyhydroxyalkanoates). Our dual-use concept aligns with the principles of a circular bioeconomy, wherein the by-products of one process become the inputs for another. By coupling plastic degradation and biomass utilization, the process can potentially be self-economically driven and minimize waste. This dual production system – obtaining both a functional enzyme and useful biomass from one cultivation – is a novel approach not demonstrated in traditional hosts like yeast or *E. coli*. Those systems generally produce a single output (the recombinant protein), whereas *C. pacifica* offers a multiple-for-one approach.

Moving forward, there are several avenues to build on this work. On the biological side, further engineering of *C. pacifica* could improve protein yield and range. For instance, constructing strains with multiple gene copies. It would also be valuable to test the secretion of other plastic-degrading enzymes, such as FASTPetase or enzymes targeting other polymers (polyurethanes, polyethylene, etc.) (47,48), to assess *C. pacifica*’s capacity as a general platform for waste-degrading biocatalysts. On the process side, scaling beyond 80 L ponds to outdoor raceway ponds or photobioreactors will be crucial for industrial adoption. Our greenhouse experiments provided controlled but realistic conditions; an outdoor trial would introduce additional variables (rainfall dilution, microbial intruders, etc.) that the extremophile *C. pacifica* might handle. The long-term stability of the transgenic trait in open environments will also need monitoring, as genetic drift or horizontal gene transfer is a consideration for any environmental GMO deployment. From an application standpoint, pairing the algae-derived enzymes with plastic waste streams will require some pretreatment of the plastics (e.g., shredding, mild heating) to improve enzyme access, as is common in enzymatic recycling schemes (49). Life-cycle assessment of the entire process would help quantify the net environmental benefit over conventional plastic disposal.

In conclusion, the expression of the PETase PHL7 in *C. pacifica* and its deployment in a pond system represent a promising convergence of synthetic biology and environmental biotechnology. We have shown that an extremophilic microalga can serve dually as a producer of potent degradative enzymes and potentially a producer of biomass feedstock, all in a single cultivation cycle. This work lays the foundation for larger-scale, sustainable algae-based platforms to tackle plastic pollution. By comparing our system with established hosts and exploring the enzymatic degradation of both standard and novel plastics, we highlight both the strengths (sustainability, low contamination risk, enzyme stability) and limitations (current yield and substrate scope) of the approach. Continued improvements in algal genetic engineering, combined with materials science innovations in creating enzyme-friendly plastics, could significantly advance the feasibility of biological recycling and bioremediation. Ultimately, integrating living systems like *C. pacifica* into waste management streams may help close the loop on plastics and contribute to a more circular, climate-friendly economy.

## Materials and Methods

### Assembly of transformation vectors

The expression vectors used in this study are derivatives of the previously described pJP32 vector (32), with full sequences provided in the Supplementary Dataset. These constructs were assembled using the pBluescript II KS+ (pBSII) backbone and the NEBuilder® HiFi DNA Assembly method (New England Biolabs, Ipswich, MA, USA). For construction of the pJP32-PHL7 vector, the PHL7 gene was codon-optimized for expression in Chlamydomonas reinhardtii and synthesized by Integrated DNA Technologies (IDT, San Diego, CA, USA). The synthetic gene was inserted into the vector backbone via HiFi DNA Assembly, following the protocol available at protocols.io (50). The vector backbone was PCR-amplified with primers containing 20 bp homology arms complementary to the ends of the PHL7 insert. All final expression vectors were designed with flanking restriction sites—XbaI at the 5′ end and KpnI at the 3′ end—to enable linearization. Restriction enzymes used throughout were obtained from New England Biolabs. Final plasmid sequences are available at ZENODO (https://zenodo.org/records/15411934), and annotated vector maps are presented in **Supplementary Figure 3**.

### Nuclear Transformation of *Chlamydomonas pacifica* and Growth Assays

Nuclear transformations were carried out using the wild-type, cell wall–containing *Chlamydomonas pacifica* strain CC-5699 (22), obtained from the Chlamydomonas Resource Center (St. Paul, MN, USA). Cultures were maintained in HSM-Acetate (HA) medium at 25 °C under constant illumination (80 μmol photons m^-2^ s^-1^) with orbital shaking at 150 rpm. Growth kinetics were assessed using a microplate-based assay adapted from a previously published protocol (51). Briefly, 160 μL aliquots from 250 mL batch cultures were sampled daily and transferred into 96-well plates, followed by absorbance measurements using an Infinite® M200 PRO plate reader (Tecan, Männedorf, Switzerland). All growth curves were based on three independent biological replicates per strain. For nuclear transformation, *C. pacifica* cultures were grown to mid-log phase (3–6 × 10^6^ cells/mL) under the conditions described above (52). Cells were harvested by centrifugation at 3000 × *g* for 10 minutes and resuspended in MAX Efficiency™ Transformation Reagent for Algae to a final density of 3–6 × 10^8^ cells/mL. Cells were then incubated on ice for 5–10 minutes with 500 ng of linearized plasmid DNA (double-digested) prior to electroporation (25). Electroporation was performed using a Gene Pulser® system (Bio-Rad) with a 4 mm gap cuvette at a field strength of 2000 V/cm and a pulse duration of 20 μs. The time constant for successful pulses was approximately 80 μs. Following electroporation, cells were transferred to 10 mL of HA medium and allowed to recover under low light at 25 °C for 18 hours with gentle agitation. Recovered cells were centrifuged, resuspended in 600 μL of fresh HA medium, and plated onto HA-agar plates supplemented with 15 μg/mL zeocin and either 0.5% or 0.75% (v/v) Impranil® DLN to enable both selection and functional screening. Plates were incubated at 25 °C under moderate light (60 μmol photons m^-2^ s^-1^) until visible colony formation was observed.

### Open Raceway Pond Preparation

Starter cultures of *C. pacifica* were initiated in 250 mL Erlenmeyer flasks containing HSM-Acetate (HA) medium supplemented with ampicillin (100 mg/mL) and the antifungal agent carbendazim (10 mg/mL) to suppress bacterial and fungal contaminants. Cultures were incubated for approximately 5 days under continuous illumination at 80 μmol photons m^-2^ s^-1^ with shaking. Once sufficient stationary phase was achieved, each culture was scaled up into 20 L carboys filled with HA medium. Carboys were cultivated under increased illumination (120 μmol photons m^-2^ s^-1^ at the surface) and aerated using air-lift mixing. After approximately 4 days, when cultures reached visually dense cell concentrations, the carboys were transported to the greenhouse field station for pond inoculation. Three 80 L open raceway ponds, housed within a greenhouse for secondary containment, were used for scale-up cultivation. Prior to use, all ponds were cleaned in place with a 10% bleach solution and thoroughly rinsed with deionized water. Each pond was then filled with 80 L of freshly prepared HSM medium, with nitrate as N source, consisting of water, salts, and trace elements, and the pH was adjusted to 10.5. Cultures from the three carboys were combined and evenly distributed across the ponds to ensure a uniform starting inoculum. Immediately following inoculation, baseline samples were collected from each pond for biomass determination, and environmental parameters—including pH and temperature—were recorded to establish the initial conditions for the experiment.

### Colony Screening and Enzyme Activity Assay

Transformants were screened for PETase activity by visual assessment of halo formation on HA agar plates supplemented with 15 *μ*g/mL zeocin and Impranil® DLN at either 0.5% or 0.75% (v/v), according to a standardized screening protocol (53). Halos were defined as clear, transparent zones surrounding colonies, indicating localized degradation of the Impranil® DLN polymer suspended in the agar matrix and thus the presence of active secreted enzyme (25).

Individual colonies exhibiting visible halos were selected and inoculated into 160 *μ*L of HA medium in Nunc™ Edge™ 96-Well Microplates (Nunclon Delta-Treated, flat-bottom; Thermo Scientific™). A total of 84 colonies were selected for screening, along with six wild-type colonies and six media-only wells as negative and blank controls, respectively. Plates were incubated under continuous illumination (60 μmol photons m^-2^ s^-1^) on a Thermo Labline Model 4625 titer plate shaker (Thermo Scientific, Iowa, USA) set to 800 rpm for five days.

After incubation, absorbance and fluorescence measurements were taken using an Infinite® M200 PRO microplate reader (Tecan, Männedorf, Switzerland), using the full set of detection parameters specified in the accompanying Data Settings file. To establish a baseline for enzymatic activity and secretion, six biological replicates of the parental wild-type strain CC-5699 were included as negative controls.

Following measurement, cultures were centrifuged at 3000 × g for 5 minutes, and the resulting supernatants were used in enzymatic activity assays. The remaining cell pellets were transferred to Impranil-containing rectangular agar plates using a sterile microplate replicator to confirm each clone’s ability to form halos under the same screening conditions.

### Plate reader settings

The Infinite® M200 PRO plate reader (Tecan, Männedorf, Switzerland) plate reader was used to measure both cell density and PHL7 enzymatic activity. Cell density was measured via chlorophyll fluorescence with an excitation of 440 nm, emission of 680 nm, and an absorbance measurement at 750 nm (22). Enzymatic activity was measured using various protocols; Enzymatic activity was also probed by using a plastic dispersion protocol with Impranil® DLN (Bayer Corporation, Germany). Similar to the Impranil/agar plates, an agarose/Impranil assay was designed which contained 0.2% (m/v) agarose and 0.25% Impranil® DLN (v/v) to keep Impranil® DLN in suspension in 96 well clear bottom plate, with readings every 5 min to measure degradation 160uL of this mixture was pipetted using a multichannel pipette into 96 well, UV-transparent, flat-bottomed plates. The enzyme’s ability to degrade post-consumer plastic was followed by taking absorbance readings at 240 nm, which were made at time 0, and after 7 days of incubation at 70 °C, in a PCR tube with PET beads with 50% crystallinity in Buffer pH8, 500 mM (54).

### Zymogram

The TGX Stain-Free™ FastCast™ Acrylamide Starter Kit (Bio-Rad Laboratories, USA) was used to prepare upright, SDS-PAGE zymogram gels containing the aforementioned Impranil PU dispersion. The acrylamide solution was mixed as per the manufacturer’s instructions, with the modification of adding 1% v/v Impranil® DLN to the solution to enable the detection of enzyme activity. This mixture was then poured into a casting frame and allowed to polymerize. Post-polymerization, the gel was placed in the electrophoresis apparatus and run under standard protein gel conditions (150V, 50 min) in SDS-Tris_Glycine buffer. The protein samples were prepared for electrophoresis by adding 4X Laemmli buffer (#1610747, Bio-Rad Laboratories, USA) without a reducing agent, and loaded directly into the gel, This allowed the run front to be visualized without entirely denaturing the enzymes. Following electrophoresis, the gel was immersed in a 100 mM Potassium Phosphate buffer solution, pH 8.0, and incubated at 37°C until transparent bands (clearing zones) appeared. This incubation step was crucial for developing clearing zones, which act as an indicator of the enzymatic degradation of the Impranil® DLN within the gel matrix. Clearing zones typically emerged within 1-2 days of incubation.

### Quantification of PET Degradation by Terephthalic Acid Release

Enzymatic degradation of PET was quantified by measuring the release of terephthalic acid (TPA), a primary hydrolysis product, via UV absorbance at 240 nm using UV-Star™ 96-well UV-transparent microplates (Greiner Bio-One). PET plastic beads were pre-washed with 1 M potassium phosphate buffer (pH 8.0) and prepared as a slurry containing 20–30% (w/v) solids. In individual PCR tubes, 100 μL of PET slurry was mixed with 100 μL of 60× concentrated enzyme-containing culture supernatant. Three biological replicates were prepared per condition.

Samples were homogenized by pipetting and briefly centrifuged at 2000 × *g* for 1 minute. A 100 μL aliquot of the supernatant was immediately transferred to a UV-transparent 96-well plate, and absorbance at 240 nm was measured using an Infinite® M200 PRO plate reader (Tecan) to establish a baseline (T0).

The remaining reaction mixtures were incubated at 70 °C for 7 days in a thermocycler with a heated lid set to 105 °C to prevent condensation. The samples were cooled to room temperature, centrifuged at 2000 × *g* for 1 minute, and 100 μL of the supernatant was transferred to fresh UV-Star™ plates for absorbance measurement at 240 nm.

A negative control using the culture supernatant from wild-type *C. pacifica* was included to account for background absorbance and non-enzymatic PET hydrolysis. PET degradation was quantified by comparing the absorbance values of experimental samples to those of the control.

### Protein Sequencing

A 50 mL culture sample from Pond 1 at day 12 was collected, and the supernatant was filtered using a .2 micron filter. The supernatant containing PHL7 enzymes was concentrated 50X using 10 kDa centrifugal filters (Amicon® Ultra, Darmstadt, Germany) to obtain a final total protein concentration of 102.6 *μ*g/mL (Qubit). The protein sample was diafiltrated with ddH_2_O, and mass spectrometry was performed at the Biomolecular and Proteomics Mass Spectrometry Facility at UC San Diego using a LUMOS Orbi-Trap. Their full protocol can be found under [“Protocols.”](https://bpmsf.ucsd.edu/training-protocols/protocols.html)

## Data Analysis

R Statistic version 4.3.2 running in the RStudio 2023.09.1+494 “Desert Sunflower” was used to import and process data, generate statistical summaries, and associated data plots. The codes used are deposited at Zenodo (https://zenodo.org/records/15411934). The data herein were collected from experiments in which pJP32PHL7 was used to transform the WT strain CC-5699. For the analyses of flask cultures, standard deviation bars represent the variation across three biological replicates of each strain.

