## Supplementary material for "Sustainable production of plastic-degrading enzymes in *Chlamydomonas pacifica*": air_temp_sensor.R

```
library(ggplot2)
library(readxl)


# Set working directory
wd <- setwd("G:/My Drive/Pesquisa/Projetos/PosDoc/Artigos Escritos/5_Plastic degrading enzyme_403/Data")

# Load the Excel file
file_path <- "G:/My Drive/Pesquisa/Projetos/PosDoc/Artigos Escritos/5_Plastic degrading enzyme_403/Data/Temp_sensor.xlsx"

# Read the MEAS data from both sheets
temperature <- read_excel(file_path)


# Remove rows with missing Temperature values
clean_data <- na.omit(temperature[, c("Time", "Temperature°C")])

# Convert 'Time' to a datetime format
clean_data$Time <- as.POSIXct(clean_data$Time, format="%Y-%m-%d %H:%M:%S")

# Plot the temperature data
ggplot(clean_data, aes(x = Time, y = `Temperature°C`, group = 1)) +
  geom_line(color = "black", size = 1, alpha = 0.7) +  # Black line
  geom_point(color = "black", size = 1, alpha = 0.5) +  # Black points
  theme_classic() +
  labs(
    title = "Temperature Sensor Data (Individual & Summary)",
    x = "Date",
    y = "Temperature (°C)"
  ) +
  theme(
    axis.text = element_text(face = "bold"),  # Bold text for axis labels
    axis.title = element_text(face = "bold"),  # Bold text for axis titles
    legend.position = "none"  # Remove legend
  ) +
  scale_x_datetime(
    date_breaks = "2 days",  # Major ticks every 2 days
    date_minor_breaks = "1 day",  # Minor ticks every day
    date_labels = "%Y-%m-%d",  # Date format in YYYY-MM-DD
    limits = c(as.POSIXct("2024-03-14"), as.POSIXct("2024-03-28"))  # Set x-axis limits
  ) +
  theme(
    axis.ticks.length = unit(0.3, "cm"),  # Major tick marks length
    axis.ticks = element_line(size = 1),  # Thicker major tick marks
    axis.ticks.minor = element_line(size = 0.5)  # Smaller internal tick marks
  )

ggsave("air_temp.svg", 
       plot = last_plot(),  # Use the most recent plot created
       device = "svg",      # Specify the SVG format
       width = 8,           # Width of the output image in inches
       height = 6)          # Height of the output image in inches
```
