## Supplementary material for "Sustainable production of plastic-degrading enzymes in *Chlamydomonas pacifica*": all sensors data.R

```
# Load necessary libraries
library(readr)
library(readxl)
library(dplyr)
library(lubridate)
library(gridExtra)

# Set working directory
wd <- setwd("G:/My Drive/Pesquisa/Projetos/PosDoc/Artigos Escritos/5_Plastic degrading enzyme_403/Data")

# Load the combined pH and temperature data
combined_df <- read_csv("pH_ponds_combined_results.csv")

# Load the light sensor data from the Excel file
light_sensor_data <- read_excel("Light_sensor.xlsx", sheet = "MEAS_1")


# Convert the `date` column to a proper datetime format
light_sensor_data <- light_sensor_data %>%
  mutate(Time = as.POSIXct(date, format = "%Y%m%d %H%M"))


# Now the time is properly formatted, you can merge it with the combined pH and temperature data
combined_df$Time <- as.POSIXct(paste(combined_df$Date, combined_df$Time), format="%m/%d/%Y %H:%M:%S")


# Merge the datasets based on the closest timestamps
merged_data <- full_join(combined_df, light_sensor_data, by = "Time")

# Save the merged dataset for future use
write_csv(merged_data, "all_data_merged_dataset.csv")


# First, round the pH and temperature data timestamps to 30-minute intervals
all_data_filtered <- merged_data %>%
  mutate(Time_rounded = floor_date(Time, unit = "30 minutes"))

# Left_join to ensure we are merging light data where available (handling nearest match)
nearest_merge <- all_data_filtered %>%
  group_by(Time_rounded) %>%
  summarise(
    mean_pH = mean(Ch1_Value, na.rm = TRUE),
    mean_temp = mean(Ch2_Value, na.rm = TRUE)
  ) %>%
  left_join(light_sensor_data %>% mutate(Time_rounded = round_date(Time, "30 minutes")),
            by = "Time_rounded")


# Step 3: Filter to keep only data from March 14 to March 28, 2024
time_filtered_data <- nearest_merge %>%
  filter(Time_rounded >= as.POSIXct("2024-03-14") & Time_rounded <= as.POSIXct("2024-03-28"))

# Step 4: Customize the x-axis breaks to every 2 days
time_breaks <- seq(from = as.POSIXct("2024-03-14"), 
                   to = as.POSIXct("2024-03-28"), 
                   by = "2 days")

# Light sensor plot (remove title, no x-axis label)
light_plot <- ggplot(time_filtered_data, aes(x = Time_rounded, y = `30-min PFD Avg`)) +
  geom_line(color = "blue") +
  scale_x_datetime(breaks = time_breaks, labels = date_format("%b %d")) +
  labs(x = NULL, y = "Light (PFD Avg)") +
  theme_minimal() +
  theme(axis.title.x = element_blank(),
        plot.title = element_blank())

# Temperature plot (remove title, no x-axis label)
temperature_plot <- ggplot(time_filtered_data, aes(x = Time_rounded, y = mean_temp)) +
  geom_line(color = "red") +
  scale_x_datetime(breaks = time_breaks, labels = date_format("%b %d")) +
  labs(x = NULL, y = "Temperature (°C)") +
  theme_minimal() +
  theme(axis.title.x = element_blank(),
        plot.title = element_blank())

# pH plot (remove title, keep x-axis label)
ph_plot <- ggplot(time_filtered_data, aes(x = Time_rounded, y = mean_pH)) +
  geom_line(color = "green") +
  scale_x_datetime(breaks = time_breaks, labels = date_format("%b %d")) +
  labs(x = "Time", y = "pH") +
  theme_minimal() +
  theme(plot.title = element_blank())


# Step 1: Open an SVG device
svg("all sensors.svg", width = 8, height = 10)  # Adjust width and height as necessary

# Step 2: Plot the grid to the SVG file
grid.arrange(light_plot, temperature_plot, ph_plot, ncol = 1)

# Step 3: Close the SVG device to save the file
dev.off()
```
