## Supplementary material for "Sustainable production of plastic-degrading enzymes in *Chlamydomonas pacifica*": light_sensor_DLI.R

```
library(ggplot2)
library(readxl)

# Set working directory
wd <- setwd("G:/My Drive/Pesquisa/Projetos/PosDoc/Artigos Escritos/5_Plastic degrading enzyme_403/Data")

# Load the Excel file
file_path <- "G:/My Drive/Pesquisa/Projetos/PosDoc/Artigos Escritos/5_Plastic degrading enzyme_403/Data/Light_sensor.xlsx"

# Read the DLI data from both sheets
dli_1 <- read_excel(file_path, sheet = "DLI_1")
dli_2 <- read_excel(file_path, sheet = "DLI_2")


# Convert Date to proper date format for Sensor 1
dli_1$Date <- as.Date(as.character(dli_1$Date), format="%Y%m%d")

# Convert Date to proper date format for Sensor 2
dli_2$Date <- as.Date(as.character(dli_2$Date), format="%Y%m%d")

# Bar plot for Sensor 1 (DLI_1)
plot1 <- ggplot(dli_1, aes(x = Date, y = `DLI (mol/(day*m*m))`)) +
  geom_bar(stat = "identity") +
  scale_x_date(date_breaks = "2 days", date_labels = "%Y-%m-%d") +
  labs(title = "DLI for Sensor 1", x = "Date", 
       y = expression("DLI Value (mol of photons/d."~m^2~")")) +
  theme_minimal() +
  theme(axis.text.x = element_text(angle = 45, hjust = 1))

# Save Sensor 1 plot as SVG
ggsave("DLI_Sensor_1.svg", plot = plot1, device = "svg")

# Bar plot for Sensor 2 (DLI_2)
plot2 <- ggplot(dli_2, aes(x = Date, y = `DLI (mol/(day*m*m))`)) +
  geom_bar(stat = "identity") +
  scale_x_date(date_breaks = "2 days", date_labels = "%Y-%m-%d") +
  labs(title = "DLI for Sensor 2", x = "Date", 
       y = expression("DLI Value (mol of photons/d."~m^2~")")) +
  theme_minimal() +
  theme(axis.text.x = element_text(angle = 45, hjust = 1))

ggsave("DLI_Sensor_2.svg", plot = plot2, device = "svg")


# Ensure matching dates and sort
dli_matched <- inner_join(dli_1, dli_2, by = "Date", suffix = c("_1", "_2"))

# Paired t-test
t_test_result <- t.test(dli_matched$DLI_1, dli_matched$DLI_2, paired = TRUE)
print(t_test_result)

# Rename columns
colnames(dli_1)[which(names(dli_1) == "DLI (mol/(day*m*m))")] <- "DLI_1"
colnames(dli_2)[which(names(dli_2) == "DLI (mol/(day*m*m))")] <- "DLI_2"

# Merge by date
dli_combined <- inner_join(dli_1, dli_2, by = "Date")

# Calculate mean and SD
dli_stats <- dli_combined %>%
  mutate(mean_DLI = rowMeans(select(., DLI_1, DLI_2), na.rm = TRUE),
         sd_DLI = apply(select(., DLI_1, DLI_2), 1, sd)) %>%
  select(Date, mean_DLI, sd_DLI)

# Plot mean with SD error bars
plot_avg_sd <- ggplot(dli_stats, aes(x = Date, y = mean_DLI)) +
  geom_bar(stat = "identity", fill = "grey", alpha = 0.8) +
  geom_errorbar(aes(ymin = mean_DLI - sd_DLI, ymax = mean_DLI + sd_DLI),
                width = 0.3, color = "black", size = 0.8) +
  scale_x_date(date_breaks = "2 days", date_labels = "%Y-%m-%d") +
  labs(title = "Average DLI Across Two Sensors",
       x = "Date",
       y = expression("Mean DLI (mol of photons/day"~m^-2~")")) +
  theme_minimal() +
  theme(axis.text.x = element_text(angle = 45, hjust = 1))

# Save plot
ggsave("DLI_Mean_SD.svg", plot = plot_avg_sd, device = "svg", width = 10, height = 6)
```
