## Supplementary material for "Sustainable production of plastic-degrading enzymes in *Chlamydomonas pacifica*": light_sensor_MEAS.R

```
library(ggplot2)
library(readxl)


# Set working directory
wd <- setwd("G:/My Drive/Pesquisa/Projetos/PosDoc/Artigos Escritos/5_Plastic degrading enzyme_403/Data")

# Load the Excel file
file_path <- "G:/My Drive/Pesquisa/Projetos/PosDoc/Artigos Escritos/5_Plastic degrading enzyme_403/Data/Light_sensor.xlsx"

# Read the MEAS data from both sheets
meas_1 <- read_excel(file_path, sheet = "MEAS_1")
meas_2 <- read_excel(file_path, sheet = "MEAS_2")


###MEAS1
# Extract Date and Time properly from the 'date' column
meas_1$Date <- as.Date(substr(meas_1$date, 1, 8), format="%Y%m%d")
meas_1$Time <- substr(meas_1$date, 10, 13)

# Combine Date and Time into POSIXct format for proper datetime handling
meas_1$DateTime <- as.POSIXct(paste(meas_1$Date, meas_1$Time), format="%Y-%m-%d %H%M")

###MEAS2
# Extract Date and Time properly from the 'date' column
meas_2$Date <- as.Date(substr(meas_2$date, 1, 8), format="%Y%m%d")
meas_2$Time <- substr(meas_2$date, 10, 13)

# Combine Date and Time into POSIXct format for proper datetime handling
meas_2$DateTime <- as.POSIXct(paste(meas_2$Date, meas_2$Time), format="%Y-%m-%d %H%M")

# Line plot for Sensor 1 (MEAS_1)
plot1 <- ggplot(meas_1, aes(x = DateTime, y = `30-min PFD Avg`)) +
  geom_line(color = "black", size = 1, alpha = 0.7) +  # Black line
  geom_point(color = "black", size = 1, alpha = 0.5) +  # Black points for visibility
  scale_x_datetime(
    date_breaks = "2 days",  # Major ticks every 2 days
    date_labels = "%Y-%m-%d %H:%M"  # Format dates with both date and time
  ) +
  labs(
    title = "Measurements for Sensor 1",
    x = "Date and Time",
    y = expression("30-min PFD Avg (unit)")
  ) +
  theme_minimal() +
  theme(
    axis.text.x = element_text(angle = 45, hjust = 1),  # Rotate x-axis labels for better readability
    axis.text = element_text(face = "bold"),  # Bold axis labels
    axis.title = element_text(face = "bold")  # Bold axis titles
  )

# Save Sensor 1 plot as SVG
ggsave("MEAS_Sensor_1.svg", plot = plot1, device = "svg")
```
