## Supplementary material for "Sustainable production of plastic-degrading enzymes in *Chlamydomonas pacifica*": pH_temp_ponds.R

```
# Set working directory
wd <- setwd("G:/My Drive/Pesquisa/Projetos/PosDoc/Artigos Escritos/5_Plastic degrading enzyme_403/Data/pH_sensor_2")


# List all CSV files in the directory
csv_files <- list.files(pattern = "*.csv")

# Read and combine all CSV files into one data frame
combined_data <- do.call(rbind, lapply(csv_files, read.csv, stringsAsFactors = FALSE))

# Save the combined data into a new CSV file in the working directory
write.csv(combined_data, "pH2_combined_results.csv", row.names = FALSE)
```
