## Supplementary material for "Sustainable production of plastic-degrading enzymes in *Chlamydomonas pacifica*": Pond_Impra.R

```
# Load libraries
library(tidyverse)
library(ggprism)

# Set the working directory
wd <- setwd("C:/Users/candi/Desktop")

# Load data
df <- read_csv("Pond_Impranil.xlsx.csv")

# Create a replicate ID (if needed)
df <- df %>%
  mutate(Replicate = row_number())  # each row is one replicate

# Pivot to long format
df_long <- df %>%
  pivot_longer(
    cols = -c(`Time [s]`, Replicate),
    names_to = "Time",
    values_to = "Value"
  ) %>%
  mutate(Time = as.numeric(Time),
         Sample = `Time [s]`) %>%
  select(-`Time [s]`)

# Compute slope per replicate
slope_df <- df_long %>%
  group_by(Sample, Replicate) %>%
  summarise(Slope = coef(lm(Value ~ Time))[["Time"]], .groups = "drop")

# Extract base sample name (e.g., R1P1 from R1P1_1)
slope_df <- slope_df %>%
  mutate(Group = case_when(
    str_detect(Sample, "^wt") ~ "wt",
    str_detect(Sample, "^R1") ~ "R1",
    str_detect(Sample, "^R2") ~ "R2",
    TRUE ~ "Other"
  ))

# Compute mean ± SD per sample (to get error bars)
sample_stats <- slope_df %>%
  group_by(Sample, Group) %>%
  summarise(
    Mean_Slope = mean(Slope),
    SD_Slope = sd(Slope),
    .groups = "drop"
  )

# Define custom colors
group_colors <- c("wt" = "#3CB043", "R1" = "#B833FF", "R2" = "#8A2BE2")

# Reorder Sample levels: wt first, then R1P1, R1P2, ..., then R2P1, ...
sample_stats <- sample_stats %>%
  mutate(Sample = factor(Sample, levels = c(
    "wt", 
    sort(unique(Sample[Group == "R1"])),
    sort(unique(Sample[Group == "R2"]))
  )))

# Plot
ggplot(sample_stats, aes(x = Sample, y = Mean_Slope, fill = Group)) +
  geom_bar(stat = "identity", color = "black") +
  geom_errorbar(aes(ymin = Mean_Slope - SD_Slope, ymax = Mean_Slope + SD_Slope), width = 0.2) +
  scale_fill_manual(values = group_colors) +
  theme_prism(base_size = 14) +
  labs(title = "Slope per Sample", y = "Slope", x = "") +
  theme(axis.text.x = element_text(angle = 45, hjust = 1)) +
  guides(fill = "none")

# ANOVA and Tukey HSD
anova_model <- aov(Slope ~ Sample, data = slope_df)
tukey_res <- TukeyHSD(anova_model)

# Extract p-values for sample pairs
tukey_letters <- multcompLetters(tukey_res$Sample[,"p adj"])
print(tukey_letters$Letters)  # View assigned letters per sample
```
