## Supplementary material for "Sustainable production of plastic-degrading enzymes in *Chlamydomonas pacifica*": production_curve.R

```
# Load necessary libraries
library(ggplot2)
library(dplyr)
library(tidyr)  
library(gridExtra)
library(ggprism)
library(grid)

# Set the working directory (update the path to your own directory)
wd <- setwd("G:/My Drive/Pesquisa/Projetos/PosDoc/Artigos Escritos/5_Plastic degrading enzyme_403/Data")

# Load the growth curve data from the provided file
growth_curve_file <- Sys.glob("20240429 Growth curve 403 wt vs 403 PHL7.csv")
growth_curve_data <- read.csv(growth_curve_file)

# Reshape the data from wide to long format
growth_curve_long <- growth_curve_data %>%
  select(-X, -X.1) %>%  # Remove unnecessary columns
  pivot_longer(cols = starts_with("X403"), 
               names_to = c("strain", "replicate"), 
               names_pattern = "X403([A-Za-z0-9]+)_([0-9]+)", 
               values_to = "abs") %>%
  mutate(strain = ifelse(strain == "Wt", "wt", "PHL7"))  # Rename strains

# Calculate mean and standard deviation for the growth curve data
growth_curve_summary <- growth_curve_long %>%
  group_by(Time..d., strain) %>%
  summarise(mean_abs = mean(abs, na.rm = TRUE), 
            sd_abs = sd(abs, na.rm = TRUE))

# Define custom colors and shapes for the strains
custom_colors <- c("wt" = "#4dac26", "PHL7" = "#d01c8b")
custom_shapes <- c("wt" = 16, "PHL7" = 17)  # Different shapes for each strain

# Plot the Growth curve (Absorbance over time) with error bars in Prism style
p1 <- ggplot(growth_curve_summary, aes(x = Time..d., y = mean_abs, color = strain, shape = strain)) +
  geom_line(size = 1.5) + 
  geom_point(size = 3) + 
  geom_errorbar(aes(ymin = mean_abs - sd_abs, ymax = mean_abs + sd_abs), width = 0.2, size = 0.8, color = "#404040") +  # Black error bars
  scale_color_manual(values = custom_colors) +  # Custom colors
  scale_shape_manual(values = custom_shapes) +  # Custom shapes
  labs(y = "OD@750nm", x = "Time (d)") +
  theme_prism(base_size = 16) +
  theme(
    axis.title.x = element_text(face = "bold"),
    axis.title.y = element_text(face = "bold"),
    legend.title = element_blank(),
    legend.position = c(0.95, 0.05),  # Legend in bottom-right corner
    legend.justification = c("right", "bottom"),  # Align to the bottom-right
    legend.background = element_rect(fill = "white", color = NA)
  )

# Save the plot
ggsave("growth_curve_plot.png", plot = p1, width = 10, height = 6, dpi = 300)
```
