## Supplementary material for "Sustainable production of plastic-degrading enzymes in *Chlamydomonas pacifica*": production_curve_pond.R

```
# Load necessary libraries
library(ggplot2)
library(dplyr)
library(lubridate)

# Set working directory
wd <- setwd("G:/My Drive/Pesquisa/Projetos/PosDoc/Artigos Escritos/5_Plastic degrading enzyme_403/Data")

# Read in the data
data <- read.csv("20240314_ponds_growth.csv")

## Simplify the column names without parentheses and spaces for easier reference
colnames(data) <- c("Elapsed_time", "DCW", "Pond")

# Convert Pond to a factor
data$Pond <- as.factor(data$Pond)

# Calculate the mean and standard deviation for each time point
data_summary <- data %>%
  group_by(Elapsed_time) %>%
  summarize(
    mean_dcw = mean(DCW),
    sd_dcw = sd(DCW)
  )

#Set the start date
start_date <- as.Date("2024-03-14")

# Create a new column in the data that represents the actual dates
data$Date <- start_date + data$Elapsed_time

# Create a new column in the summary data to represent the actual dates
data_summary$Date <- start_date + data_summary$Elapsed_time

# Now plot with dates on the x-axis
  ggplot() +
    geom_line(data = data, aes(x = Date, y = DCW, group = Pond, color = Pond), alpha = 0.5) +  # Individual ponds
    geom_point(data = data, aes(x = Date, y = DCW, group = Pond, color = Pond), alpha = 0.5) +  # Points for individual ponds
    geom_line(data = data_summary, aes(x = Date, y = mean_dcw), color = "black", size = 1) +  # Average line in black
    geom_ribbon(data = data_summary, aes(x = Date, ymin = mean_dcw - sd_dcw, ymax = mean_dcw + sd_dcw), fill = "grey", alpha = 0.2) +  # SD shading
    theme_classic() +
    labs(
      title = "Pond Growth Over Time (Individual & Summary)",
      x = "Date",
      y = "DCW (mg/L)"
    ) +
    theme(
      axis.text = element_text(face = "bold"),  # Bold text for axis labels
      axis.title = element_text(face = "bold"),  # Bold text for axis titles
      legend.position = c(0.95, 0.05),  # Bottom-right inside the graph
      legend.justification = c("right", "bottom"),
      legend.background = element_rect(fill = "white", color = "black")
    ) +
    scale_x_date(
      date_breaks = "2 days",  # Major ticks every 2 days
      date_minor_breaks = "1 day",  # Minor ticks every day
      date_labels = "%Y-%m-%d",  # Date format in YYYY-MM-DD
      limits = c(as.Date("2024-03-14"), as.Date("2024-03-28"))  # Set the x-axis limits to span from 2024-03-14 to 2024-03-28
    ) +
    theme(
      axis.ticks.length = unit(0.3, "cm"),  # Major tick marks length
      axis.ticks = element_line(size = 1),  # Thicker major tick marks
      axis.ticks.minor = element_line(size = 0.5)  # Smaller internal tick marks
    )

# Save the plot as an SVG
ggsave("pond_growth_plot.svg", 
       plot = last_plot(),  # Use the most recent plot created
       device = "svg",      # Specify the SVG format
       width = 8,           # Width of the output image in inches
       height = 6)          # Height of the output image in inches
```
