## Supplementary material for "Sustainable production of plastic-degrading enzymes in *Chlamydomonas pacifica*": tpa_activity.R

```
# Load necessary libraries
library(ggplot2)
library(ggsignif)

# Set the working directory (update the path to your own directory)
setwd("G:/My Drive/Pesquisa/Projetos/PosDoc/Artigos Escritos/5_Plastic degrading enzyme_403/Data")

# Load the CSV file
tpa_activity_data <- read.csv("20240429 TPA activity from growth curve 403 wt vs 403 PHL7.csv")

# Calculate the mean and standard deviation for each strain
tpa_activity_summary <- aggregate(delta_bsorbance ~ strain, data = tpa_activity_data, 
                                  FUN = function(x) c(mean = mean(x), sd = sd(x)))

# Split the aggregated data into separate columns for mean and standard deviation
tpa_activity_summary <- data.frame(
  strain = tpa_activity_summary$strain,
  mean_abs = tpa_activity_summary$delta_bsorbance[, "mean"],
  sd_abs = tpa_activity_summary$delta_bsorbance[, "sd"]
)

# Ensure 'wt' is displayed first
tpa_activity_summary$strain <- factor(tpa_activity_summary$strain, levels = c("wt", "PHL7"))

# Perform ANOVA analysis
anova_result <- aov(delta_bsorbance ~ strain, data = tpa_activity_data)
anova_summary <- summary(anova_result)

# Check the p-value
p_value <- anova_summary[[1]]["strain", "Pr(>F)"]

# Define custom colors for the strains
custom_colors <- c("wt" = "#4dac26", "PHL7" = "#d01c8b")

# Create the bar plot with error bars and a black border around the bars
p1 <- ggplot(tpa_activity_summary, aes(x = strain, y = mean_abs, fill = strain)) +
  geom_bar(stat = "identity", color = "black", width = 0.7) +  # Black border around the bars
  geom_errorbar(aes(ymin = mean_abs - sd_abs, ymax = mean_abs + sd_abs), width = 0.2, size = 0.8) +  # Error bars
  scale_fill_manual(values = custom_colors) +  # Custom colors
  labs(y = "Delta Absorbance", x = "Strain") +
  theme_minimal(base_size = 16) +
  theme(
    axis.title.x = element_text(face = "bold"),
    axis.title.y = element_text(face = "bold"),
    legend.position = "none"  # Remove legend
  )

# Add a significance marker if p-value is less than 0.05
if (p_value < 0.05) {
  p1 <- p1 + geom_signif(comparisons = list(c("wt", "PHL7")),
                         annotations = "*",
                         y_position = max(tpa_activity_summary$mean_abs) + 0.1,  # Adjust position of asterisk
                         tip_length = 0.01)
}

# Display the plot
print(p1)

# Save the plot
ggsave("tpa_activity.png", plot = p1, width = 10, height = 6, dpi = 300)

# Print the ANOVA result
print(anova_summary)
```
