## Supplementary material for "Sustainable production of plastic-degrading enzymes in *Chlamydomonas pacifica*": TPA_standard Curve.R

```
# Load necessary libraries
library(ggplot2)
library(dplyr)

# Raw data
data <- data.frame(
  Concentration_mM = c(
    rep(100, 3), rep(20, 3), rep(4, 3), rep(0.8, 3), rep(0.16, 3), rep(0, 3)
  ),
  OD = c(
    0.9929999709, 1.016000032, 1.033400059,
    0.2714999914, 0.2788000107, 0.2804999948,
    0.1254000068, 0.1264999956, 0.126000002,
    0.09560000151, 0.09560000151, 0.09309999645,
    0.09130000323, 0.0896999985, 0.08820000291,
    0.08110000193, 0.08160000294, 0.08049999923
  )
)

# Calculate mean OD per concentration
summary_data <- data %>%
  group_by(Concentration_mM) %>%
  summarise(
    mean_OD = mean(OD),
    sd_OD = sd(OD)
  )

# Subtract blank (0 mM) mean from all other means
blank_mean <- summary_data %>% filter(Concentration_mM == 0) %>% pull(mean_OD)
summary_data <- summary_data %>%
  mutate(corrected_OD = mean_OD - blank_mean)

# Linear regression (excluding 0 mM point)
fit <- lm(corrected_OD ~ Concentration_mM, data = summary_data %>% filter(Concentration_mM != 0))
eq <- paste0("y = ", round(coef(fit)[2], 4), "x + ", round(coef(fit)[1], 4))
r2 <- round(summary(fit)$r.squared, 4)

# GraphPad Prism-like theme
prism_theme <- theme_minimal(base_size = 14) +
  theme(
    axis.title = element_text(face = "bold"),
    axis.text = element_text(color = "black"),
    panel.grid.major = element_line(size = 0.5, linetype = 'dashed', color = 'gray'),
    panel.grid.minor = element_blank(),
    plot.title = element_text(face = "bold", hjust = 0.5)
  )

# Plot
ggplot(summary_data %>% filter(Concentration_mM != 0), aes(x = Concentration_mM, y = corrected_OD)) +
  geom_point(size = 3) +
  geom_errorbar(aes(ymin = corrected_OD - sd_OD, ymax = corrected_OD + sd_OD), width = 1) +
  geom_smooth(method = "lm", se = FALSE, color = "blue", size = 1) +
  geom_text(x = max(summary_data$Concentration_mM)*0.5, y = max(summary_data$corrected_OD)*0.9,
            label = paste(eq, "\nR² =", r2), size = 4, hjust = 0) +
  labs(
    title = "TPA Standard Curve",
    x = "Concentration (mM)",
    y = "Corrected Absorbance (OD)"
  ) +
  prism_theme
```
