## Supplementary figures and images for "Sustainable production of plastic-degrading enzymes in *Chlamydomonas pacifica*"

### 20240417 403 PHL7 PXL_20240417_162804453.jpg

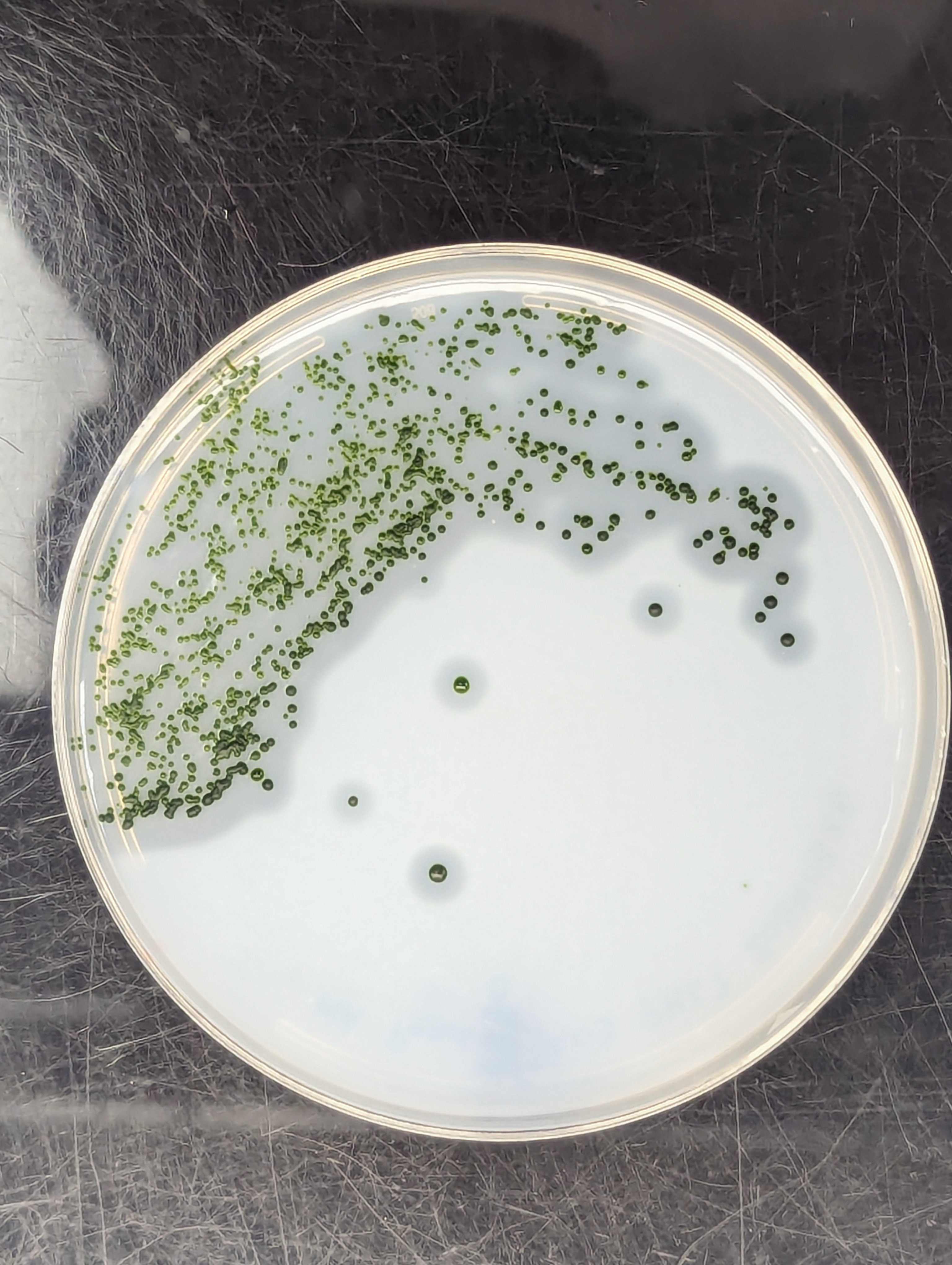

### 20240509 Zymogram Wt LA PHL7 growth curve Lab Mayfield lab 2024-05-09 12h35m07s(Amido Black) copy.tif

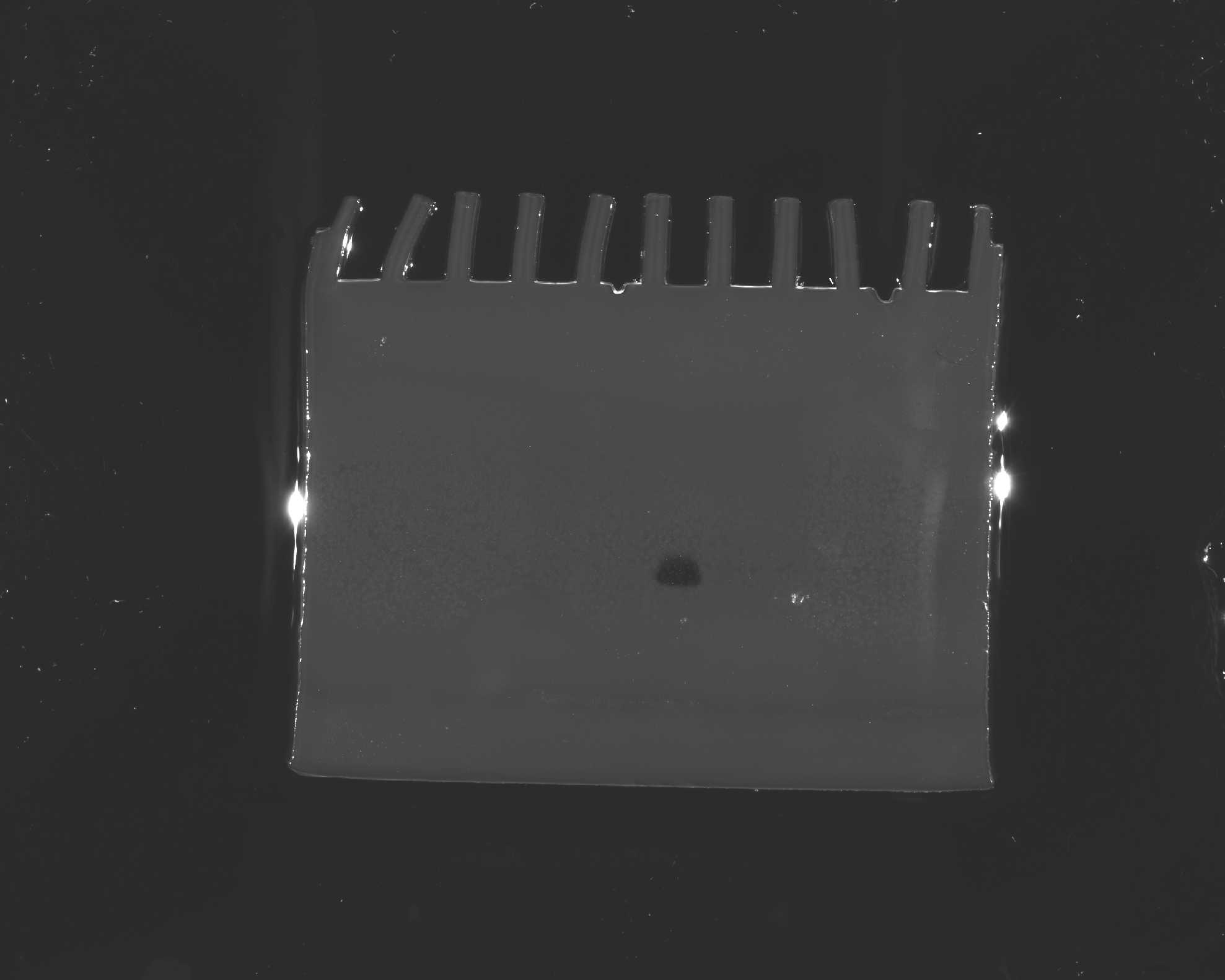

### 20240509 Zymogram Wt LA PHL7 growth curve Mayfield lab 2024-05-08 12h49m03s(Stain Free Gel).raw16.tif

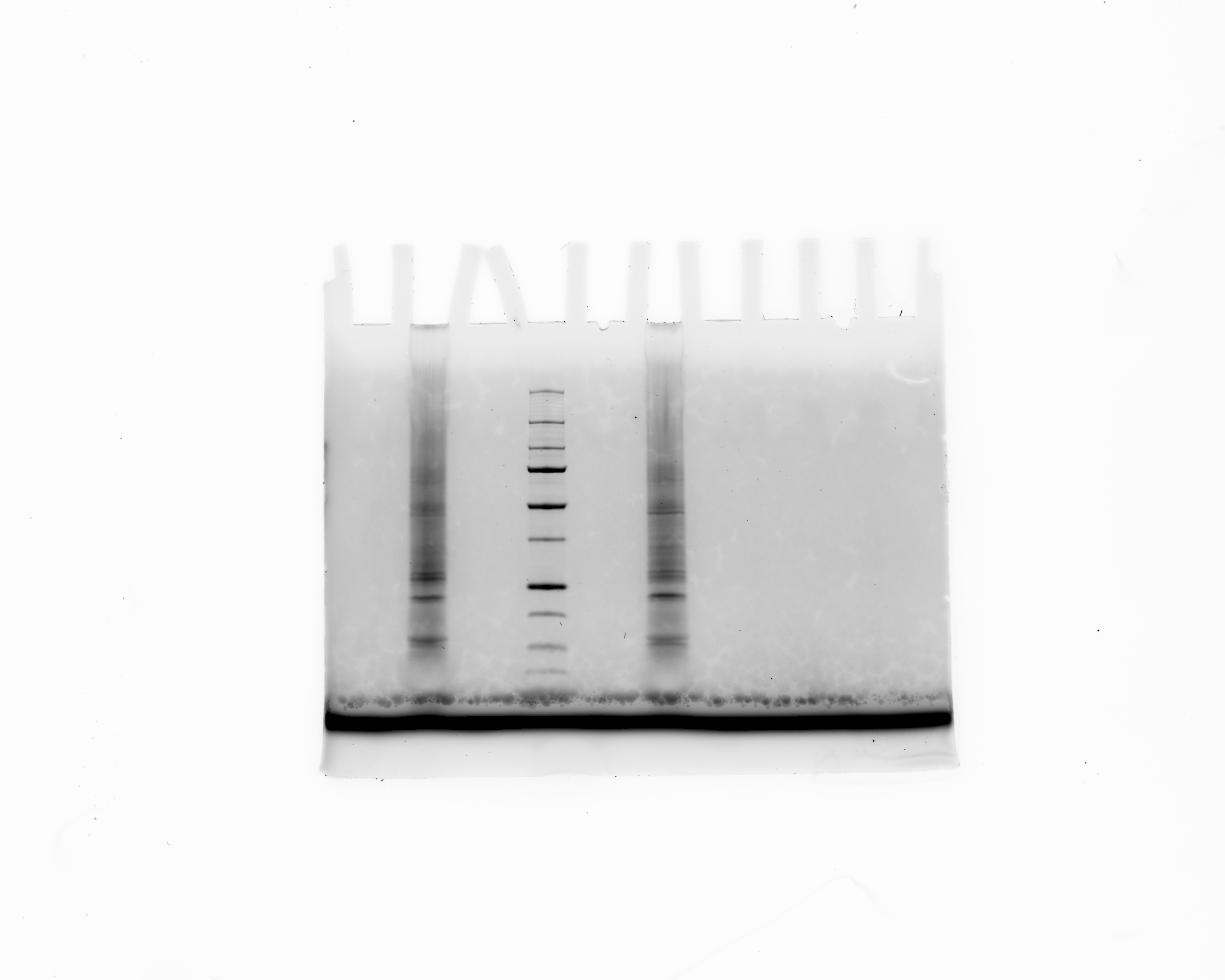

### growth_curve_plot.png

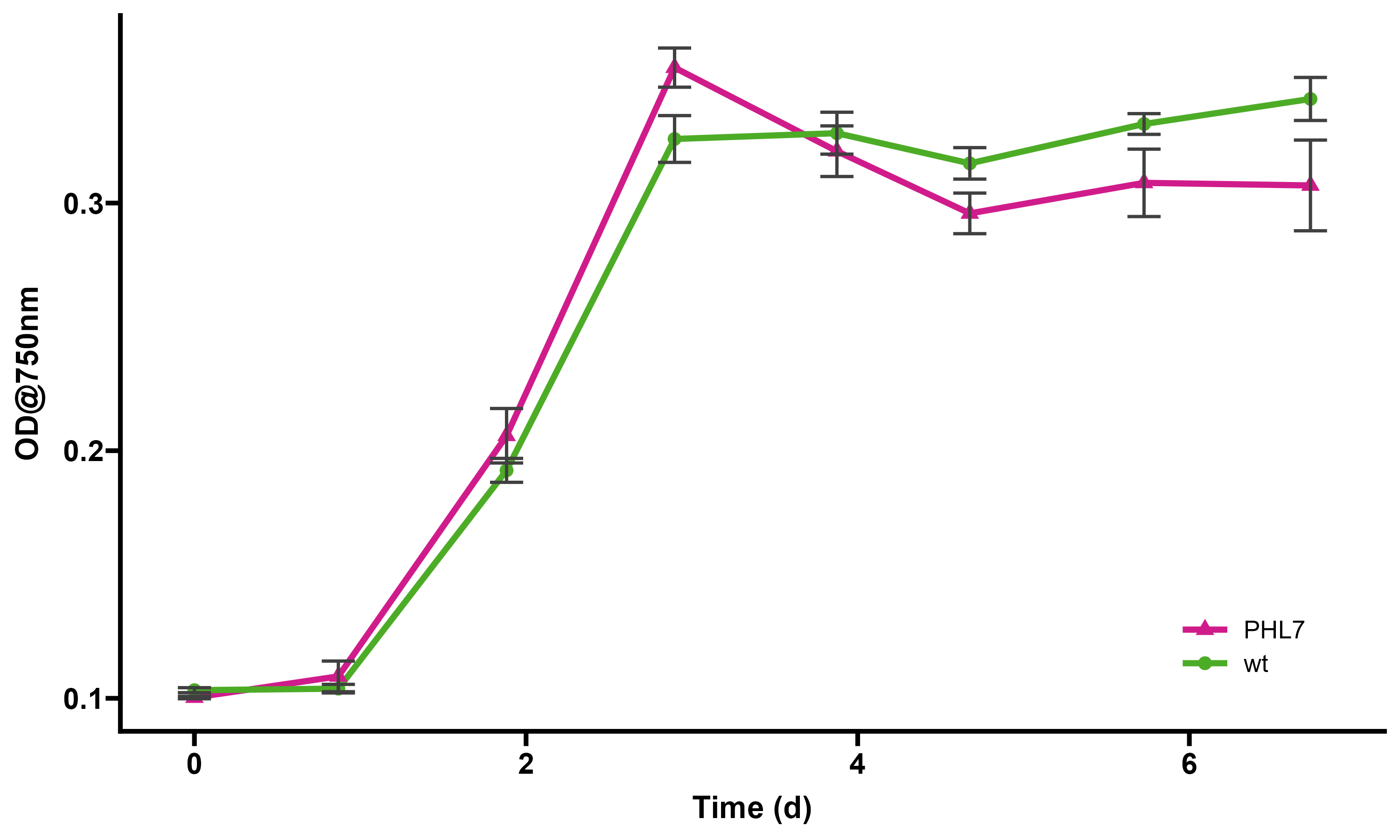

### image2894.png

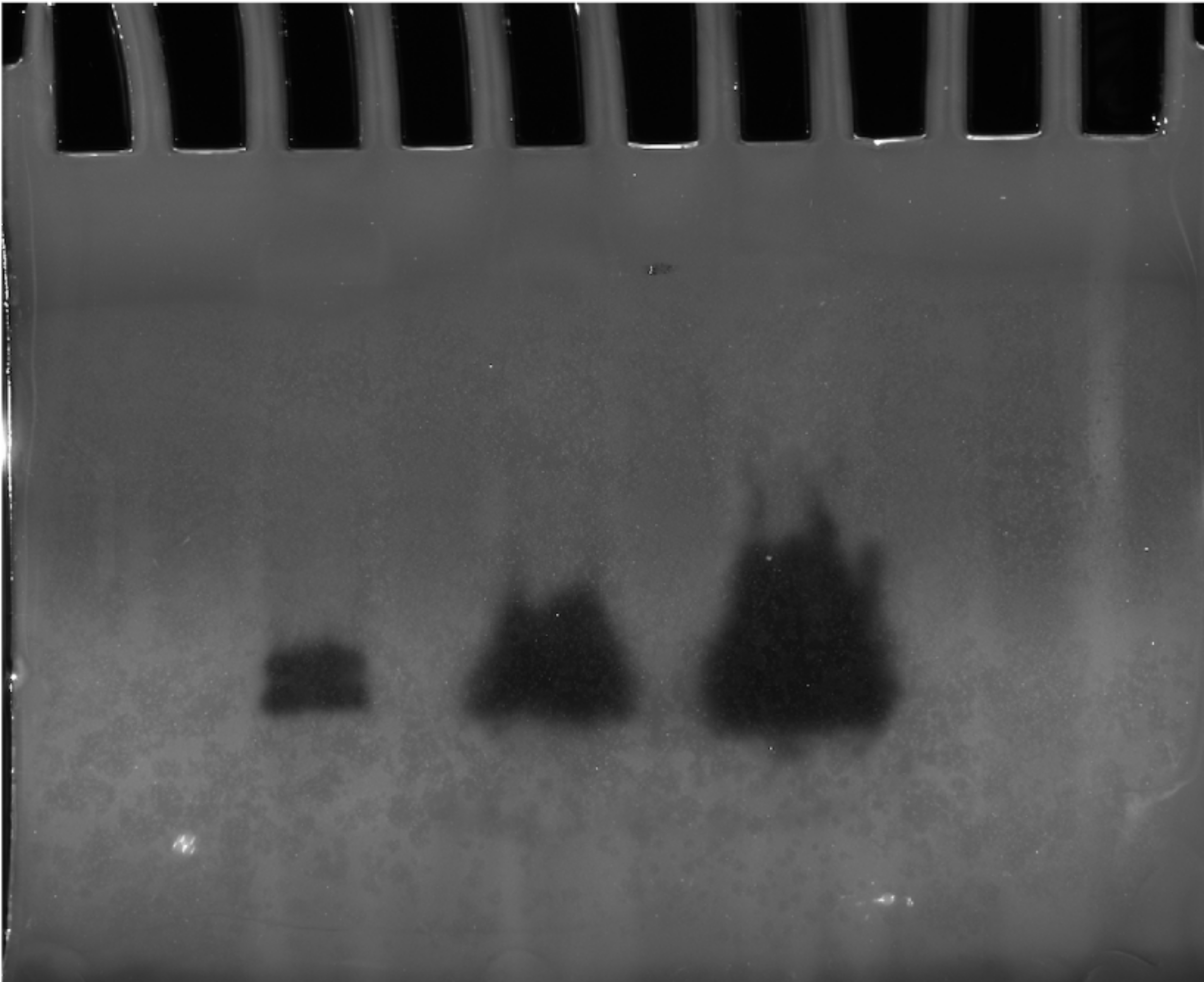

### image2906.png

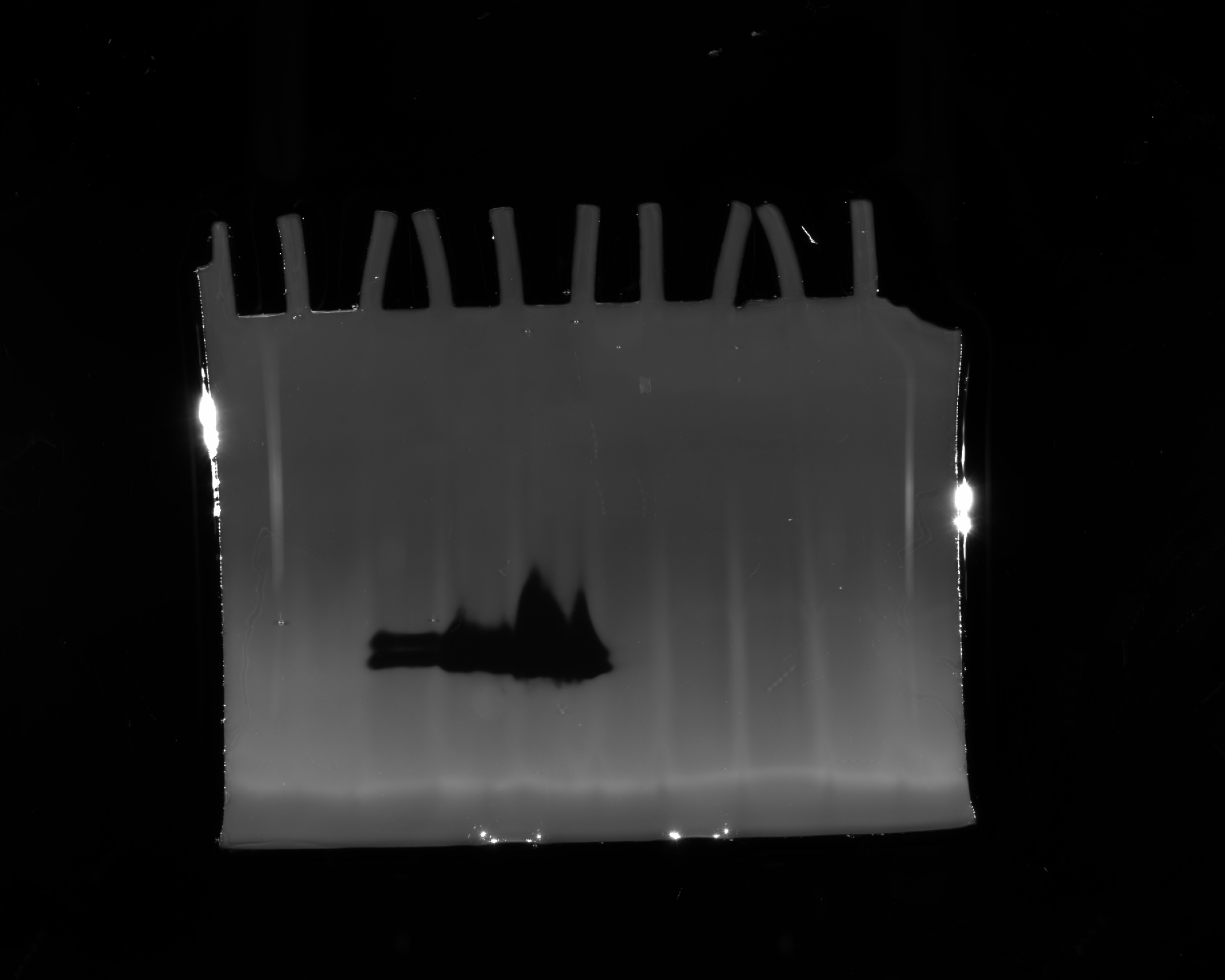

### PHL7_WT_plates_1.JPG

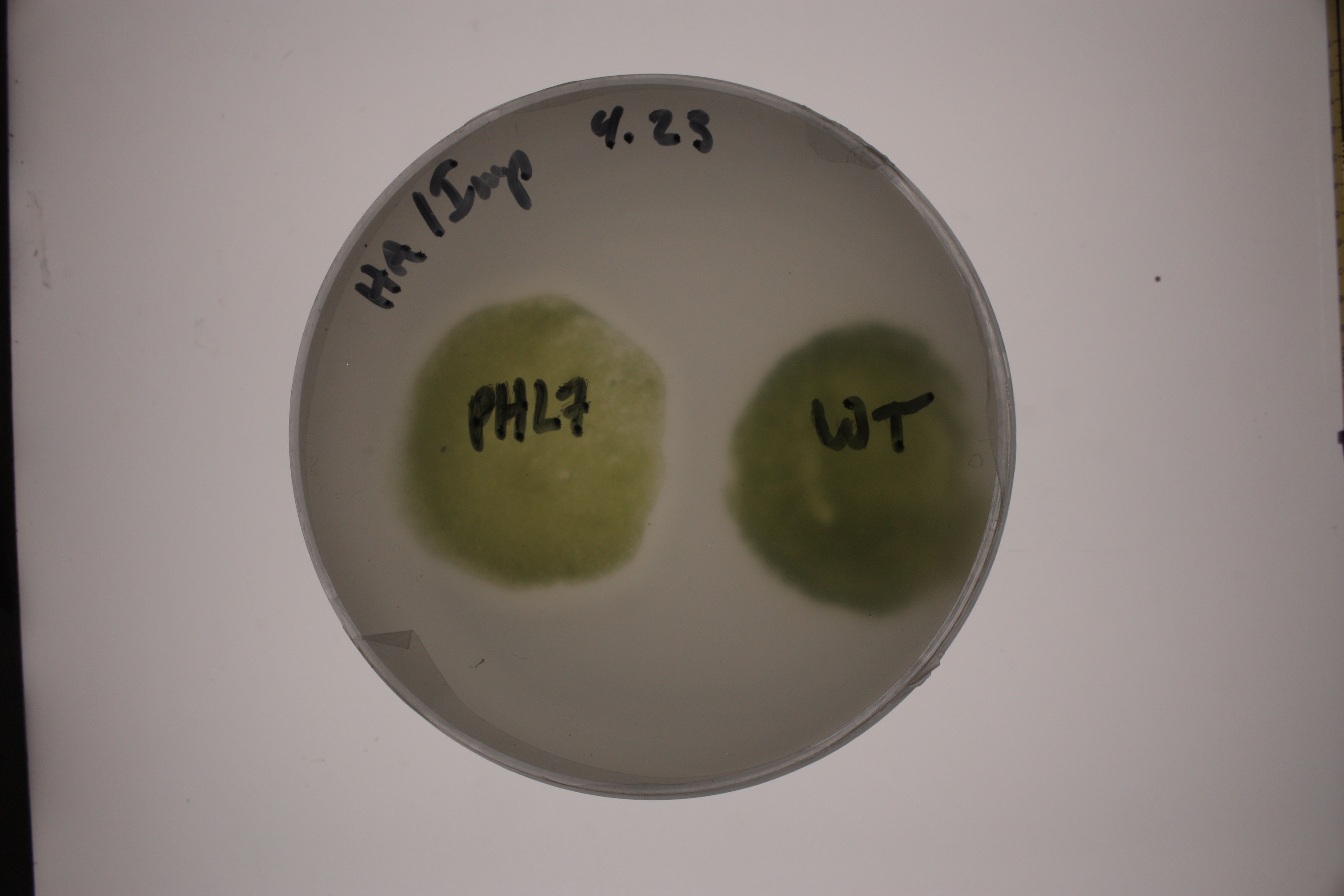

### pJP32PHL7 Map.png

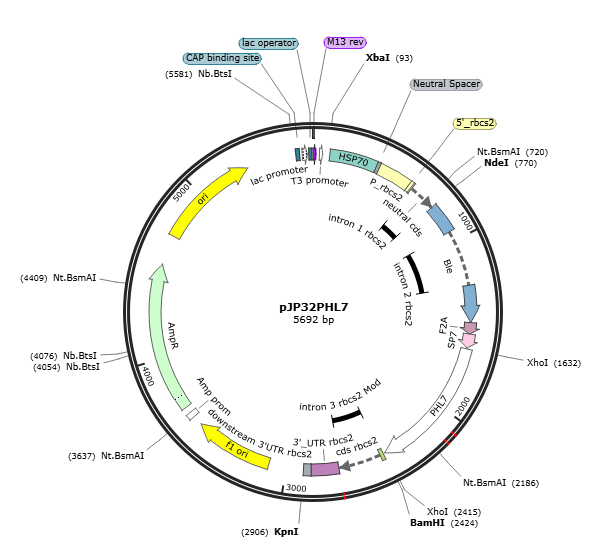

### tpa_activity_prism_style.png

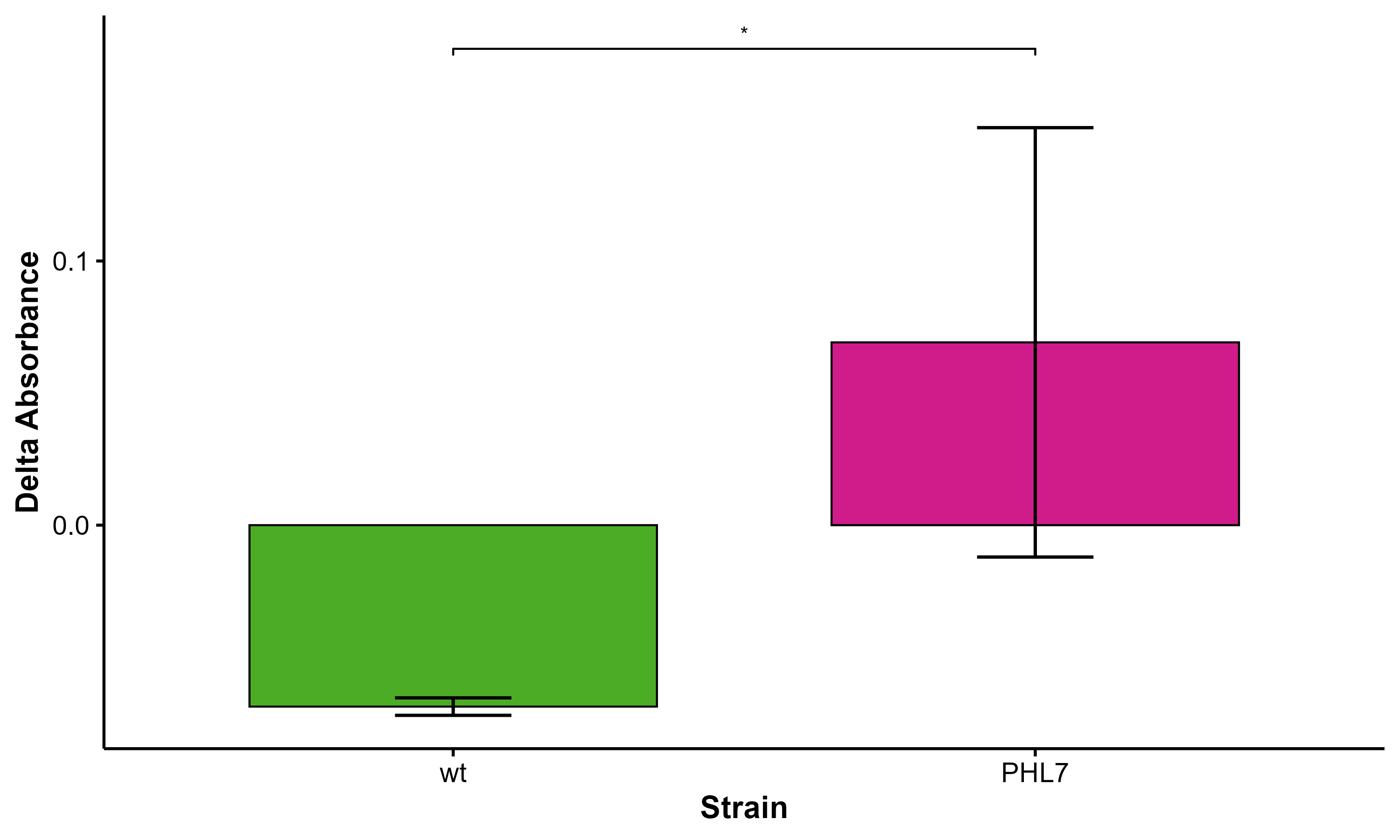
